# Functional genomic screening in *Komagataella phaffii* enabled by high-activity CRISPR-Cas9 library

**DOI:** 10.1101/2024.02.08.579509

**Authors:** Aida Tafrishi, Varun Trivedi, Zenan Xing, Mengwan Li, Ritesh Mewalal, Sean Culter, Ian Blaby, Ian Wheeldon

## Abstract

CRISPR-based high-throughput genome-wide loss-of-function screens are a valuable approach to functional genetics and strain engineering. The yeast *Komagataella phaffii* is a host of particular interest in the biopharmaceutical industry and as a metabolic engineering host for proteins and metabolites. Here, we design and validate a highly active 6-fold coverage genome-wide sgRNA library for this biotechnologically important yeast containing 30,848 active sgRNAs targeting over 99% of its coding sequences. Conducting fitness screens in the absence of functional non-homologous end joining (NHEJ), the dominant DNA repair mechanism in *K. phaffii*, provides a quantitative means to assess the activity of each sgRNA in the library. This approach allows for the experimental validation of each guide’s targeting activity, leading to more precise screening outcomes. We used this approach to conduct growth screens with glucose as the sole carbon source and identify essential genes. Comparative analysis of the called gene sets identified a core set of *K. phaffii* essential genes, many of which relate to protein production, secretion, and glycosylation. The high activity, genome-wide CRISPR library developed here enables functional genomic screening in *K. phaffii*, applied here to gene essentiality classification, and promises to enable other genetic screens.

**Highlights:**

- Designed and validated a high activity genome-wide CRISPR-Cas9 library for *K. phaffii*
- Disabling NHEJ DNA repair enables the generation of genome-wide guide activity profiles
- Activity-corrected fitness screens identify a high confidence set of essential genes in *K. phaffii*
- Protein production, secretion, and glycosylation pathways are essential in *K. phaffii* but not in other yeasts

## 1.0 Introduction

The methylotrophic yeast *Komagataella phaffii*, formerly known as *Pichia pastoris*, is commonly referred to as the “biotech yeast” because of its widespread adoption within the pharmaceutical and biotechnology industry (Ahmad et al., 2014; Bernauer et al., 2020; Daly and Hearn, 2005; Gasser et al., 2013). This microorganism has emerged as an important recombinant protein production host because it is able to grow to high cell densities as it favors respiratory growth compared to fermentative yeasts, secretes significant levels of heterologous protein in the media saving time and cost for downstream purification processes, has a strong alcohol oxidase I (*AOX1*) promoter facilitating controlled expression of recombinant genes, is able to perform post-translational modifications similar to higher eukaryotes, can assimilate a variety of carbon sources including methanol, and is a generally faster, easier and cost-efficient expression host compared to mammalian cell lines (Ata et al., 2021; Love et al., 2018).

Constructing advanced microbial cell factories requires the development of efficient genetic engineering tools. Previous efforts in *K. phaffii* engineering include the development of integrative gene expression systems through the use of homologous recombination (HR) (Cai et al., 2021; Weninger et al., 2016), the design of episomal gene expression vectors (Cregg et al., 1985), and the standardization of variable strength promoters (Cai et al., 2021). CRISPR-Cas9 has become the preferred engineering method allowing for precise, targeted, and relatively rapid genetic modifications (Dalvie et al., 2020; Liu et al., 2015; Löbs et al., 2017a, 2017b). Using CRISPR-Cas9 with pooled single guide RNA (sgRNA) libraries, allowing for genome-wide screens, has been used as a high-throughput method to analyze gene functions, assign genotypes to phenotypes, and identify essential genes (Dong et al., 2022; Lupish et al., 2022; Ramesh et al., 2023; Shalem et al., 2014; Trivedi et al., 2023). Other functional genomic tools include random chemical or transposon mutagenesis. While these methods have been used successfully in various applications (Guo et al., 2013; Kim et al., 2010; Michel et al., 2017; Patterson et al., 2018; Zhu et al., 2018), they can be limited by the random nature of the resulting mutants, which is biased towards longer genes (Wang et al., 2018). CRISPR approaches use targeted mutagenesis, ultimately producing a more diverse mutant pool and more accurate screening outcomes (Morgens et al., 2016; Ramesh et al., 2023; Schwartz et al., 2019).

One of the challenges with genome-wide CRISPR screens, particularly in non-conventional species, is accurate guide activity predictions. While a number of CRISPR guide activity predictors have been developed (Zhang et al., 2023), they are most often trained on a selected number of model species (e.g., *Escherichia coli*, *Saccharomyces cerevisiae* or mammalian cell lines) and the ability to predict active guides in other species is not well established (Baisya et al., 2022; Moreb and Lynch, 2021). A solution to this problem is to design multiple guides to target a single gene, thus biasing the library toward at least one active guide per gene. This redundancy in guide design, however, introduces complexities in downstream analysis and dramatically increases library size, which can be problematic if efficient transformation protocols are not available for the host of interest. We have addressed this problem by developing an experimental approach to generate genome-wide CRISPR activity profiles that can be used in combination with functional screens to improve screening outcomes (Ramesh et al., 2023; Schwartz et al., 2019). The basic principle is to deactivate the native dominant DNA repair mechanism, typically non-homologous end joining (NHEJ) in non-conventional yeasts such as *K. phaffii* (Bernauer et al., 2020), and conduct growth screens in the absence of DNA repair. Such screens provide an indirect measure of guide activity as any double stranded break in the genome leads to cell death or a dramatic reduction in cell fitness. The guide activity profiles can be incorporated into the screening analysis pipeline by analytically removing inactive or poorly active guides, thus improving screen accuracy (Ramesh et al., 2023).

Here, we design, validate, and deploy a 6-fold coverage, high-activity pooled CRISPR-Cas9 sgRNA library targeting over 99% of the protein-coding sequences in *Komagataella phaffii* GS115. By disabling NHEJ via functional disruption of *KU70*, we first quantify the activity of the library. This guide activity data is used to correct the outcomes of fitness screens and accurately identify essential genes with glucose as the sole carbon source. Analysis of the essential genes revealed a set of essential genes common across a collection of industrially-relevant biochemical production hosts and model yeasts, and others that are unique to *K. phaffii*. Identification of essential genes contributes to the overall understanding of *K. phaffii* genetics and enhances gene annotation that will help metabolic engineers create optimized *K. phaffii* production strains. The CRISPR screens used to generate this new data opens new functional genetic screening capabilities for the biotech yeast and promises to enable rapid metabolic engineering workflows.

## 2.0 Results and Discussion

### 2.1 Pooled sgRNA library enables functional genetic screening in *K. phaffii*

Pooled sgRNA libraries enable forward genetic screens. When transformed into a Cas9 expressing strain, each cell expresses a single sgRNA targeting a gene disruption; outgrowth of the transformants creates a pool of mutant cells with varying phenotypes. The fitness effects due to each sgRNA are quantified by determining a fitness score (FS), the log_2_ ratio of th normalized abundance of the sgRNA in sample to that of a control strain (**Figure 1a**). Similarly, a cutting score (CS) can be determined for each sgRNA by comparing the normalized abundance of guides in a NHEJ deficient strain to a control strain absent of Cas9. Since no DNA repair template is provided and the cells lack NHEJ, a double-stranded break in the genome results in cell death or a dramatic reduction in cell fitness, thus allowing us to quantify Cas9 activity for a given sgRNA. FS and CS profiles for *K. phaffii* GS115 over a six day period, including one subculture at day 3, are shown in **Figure 1b** and **Supplementary Data 1**. Fitness effects are evident after three days of growth (the first time the cultures reached confluency) and are more pronounced after subculturing the population and allowing for additional outgrowth. Notably, non-targeting controls consistently exhibit low CS and high FS values across both time points, indicating their inactivity and negligible impact on cell fitness. In contrast, targeting sgRNAs exhibit a range of CS values and have a broad effect on cell fitness.

**Figure 1.**
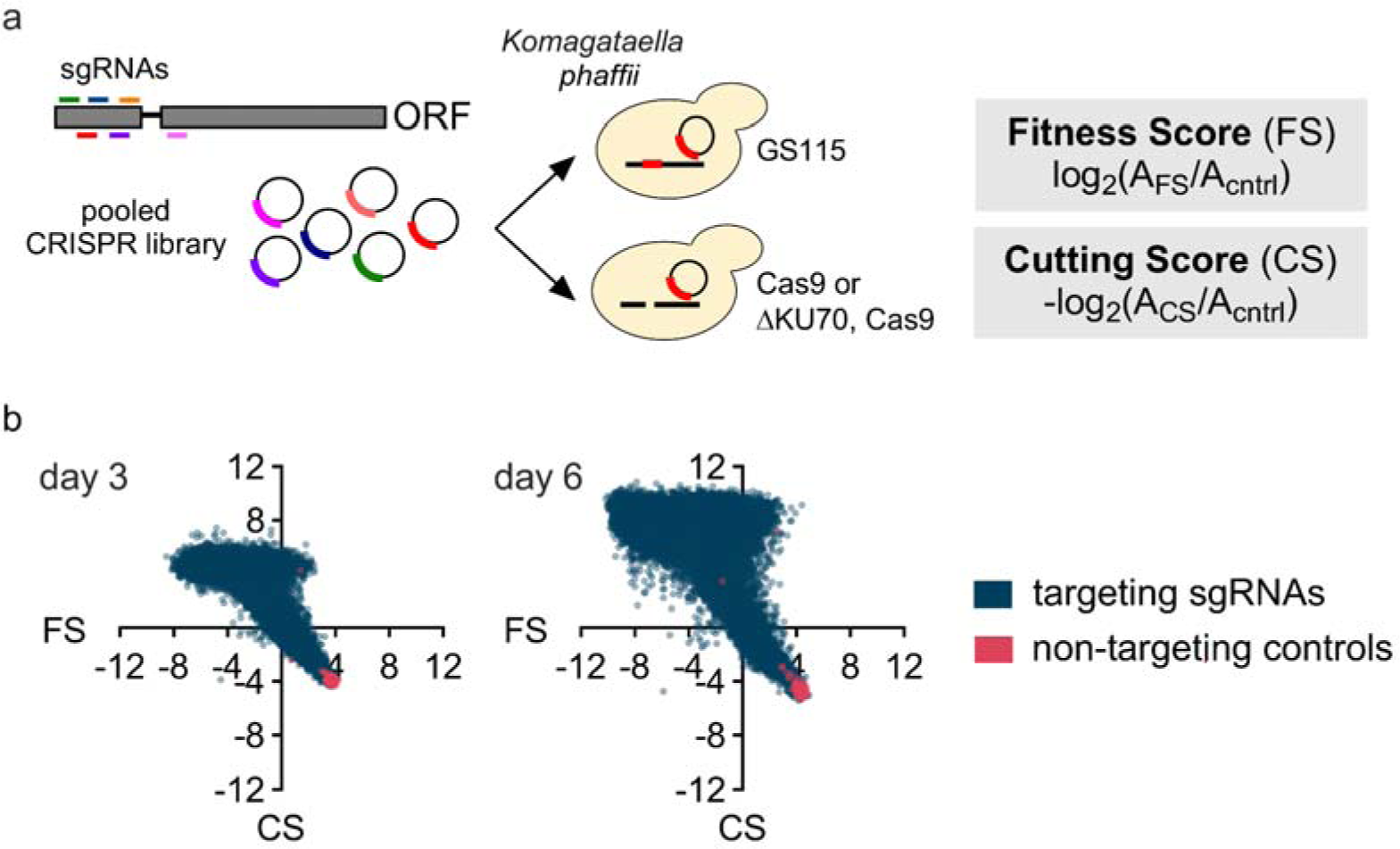
Genome-wide CRISPR-Cas9 single guide RNA (sgRNA) functional genetic screens in *K. phaffii*. **a)** Fitness and cutting score screens. *Komagataella phaffii* GS115 strain was used as the base strain for all experiments. GS115 *his4::CAS9* and GS115 *his4::CAS9* Δ*KU70* strains were used for fitness score (FS) and cutting score (CS) experiments, respectively. GS115 and GS115 Δ*KU70* were used as the control strains for the FS and CS screens. A genome-wide sgRNA library was designed to target the first 300 bp of each expressed gene. The 6-fold coverage library was transformed into each strain and growth screens were performed to determine CS and FS for each sgRNA. **b)** Scatter plots of the CS and FS values generated on day 3 and 6 of the screens. Data points represent the average FS and CS values for triplicate experiments; each replicate was created with an independent library transformation.

### 2.2 *In silico* sgRNA design produces a highly active guide library

We designed a 6-fold genome-wide sgRNA library targeting 5,309 protein coding sequence (CDS) and 120 tRNAs in *K. phaffii* GS115 (Figure 2a and **Supplementary Data 2**). The initial library included 169,034 sgRNAs targeting the first 300 bp of each CDS and tRNAs (**Supplementary Data 3**). Using a combined metric that accounted for the predicted activity of each guide and the uniqueness of each guide sequence, this large pool of guides was reduced to 31,634, including the top six ranked guides for each gene in the genome. An additional 350 non-targeting sgRNAs (randomly generated sequences with no homology to the GS115 genome, **Supplementary Data 4**) were added to the library for a total of 31,984 sgRNAs targeting 99.68% of the CDSs. Seventeen CDSs were excluded from the library due to the lack of unique guides (**Supplementary Table 1**).

**Figure 2.**
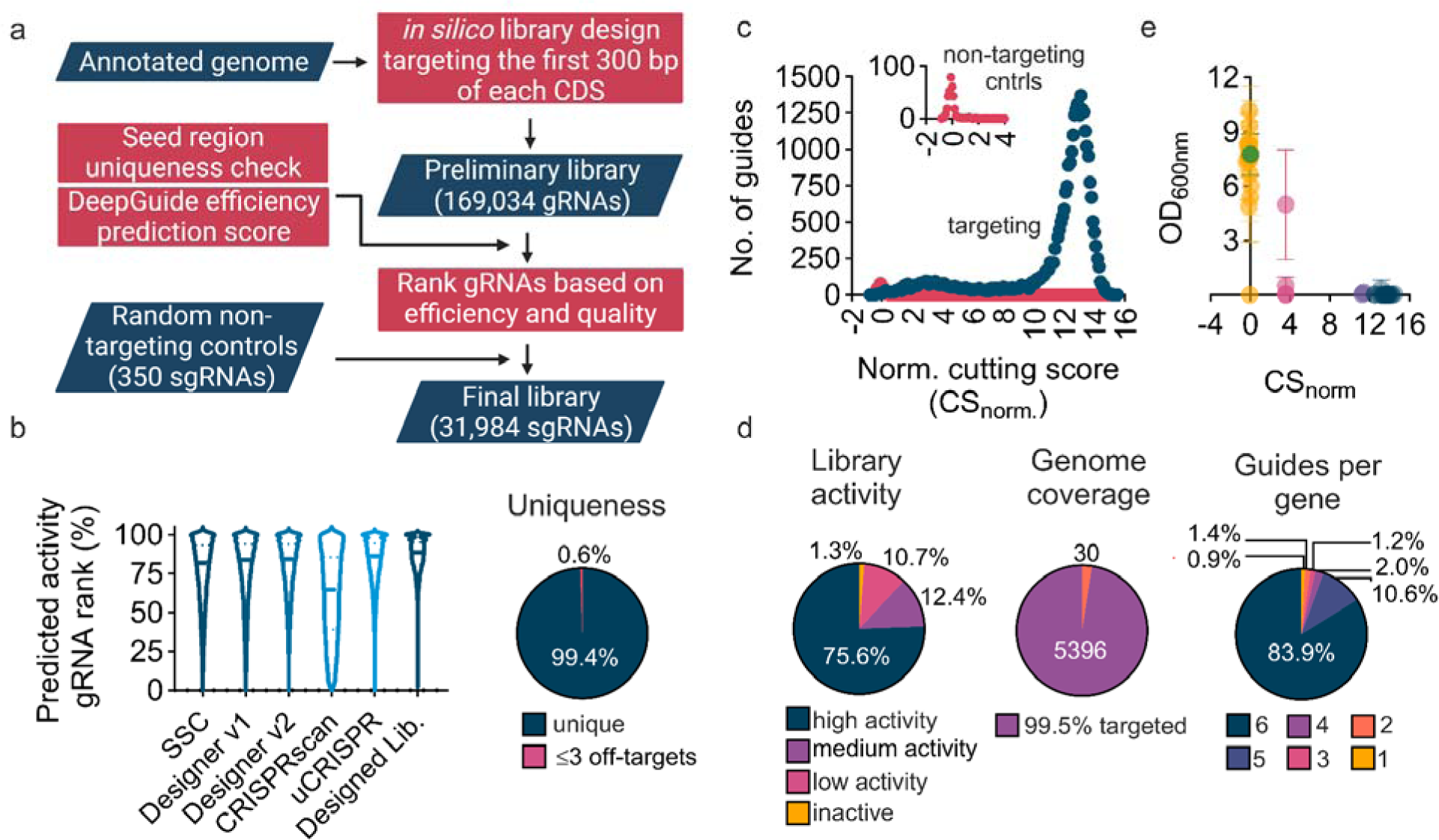
Genome-wide library design, genome-wide CS profile and validation. **a)** Schematic representation of the genome-wide sgRNA library design workflow. CHOPCHOP v3 and custom python scripts were used to identify all sgRNAs targeting the first 300 bp of each coding sequence (CDS) and tRNA genes. A series of guide activity prediction methods (five used by CHOPCHOP v3 plus DeepGuide (Baisya et al., 2022)) and a quality score (uniqueness and self-complementarity) were used to identify the best six sgRNAs targeting each gene. The final library consisted of 31,634 genome-targeting sgRNAs and 350 non-targeting controls. **b)** Criteria for choosing the best six sgRNAs for the final library. The violin plots show the ranked activity of guides as predicted by the five algorithms used by CHOPCHOP v3. 99.4% of the sgRNAs in the library are unique (see methods for uniqueness criteria) and only 0.6% of the library consists of sgRNAs with up to 3 off-targets and up to 3 mismatches (**Supplementary Table 7**). **c)** CS distribution on day 6. The CS for each sgRNA is normalized to the average CS of non-targeting controls. The presented CS values are the mean of three biological replicates. **d)** CS downstream analysis. 98.7% of the library consists of high (CS_norm_ > 11.46), medium (6.90 <CS_norm_< 11.46), and low activity (1.36 <CS_norm_< 6.90) sgRNAs. 392 sgRNAs were identified as inactive (CS_norm_ < 1.36). Active sgRNAs target 5,396 genes in the GS115 genome, with only 30 genes not covered in the library. 83% of genes were targeted with 6 active sgRNAs. **e)** CS validation. 24 active (including highly active (dark blue), medium (purple), and low (magenta) activity) and 16 inactive (yellow) sgRNAs were chosen for validation experiments. sgRNAs were expressed in GS115 *his4*::*CAS9* Δ*KU70*. Transformants with inactive sgRNAs showed growth similar to a control (green) strain, whereas cells transformed with active sgRNAs showed no or limited growth compared to control (3-day culture in SD-H, 2% glucose, 30°C, 225 rpm). Data points and error bars represent the average of three biological replicates and one standard deviation, respectively.

Guides chosen to be in the final library are highly ranked by all activity predictors and over 99% have unique seed sequences (the 12 bp upstream of the PAM sequence) with no predicted off-target effects (Figure 2b, see “Materials and Methods” for more details). We focused our uniqueness criteria on the seed sequence because off-target effects have been shown to be more prominent with mismatches outside of the seed region and seed uniqueness is critical to on-target Cas9 effectiveness (Cong et al., 2013; Hsu et al., 2013; Jiang et al., 2013; Jinek et al., 2012; Labun et al., 2019).

Using the designed library, we conducted a growth screen with cells containing disabled NHEJ to generate a CS profile across the genome (Figure 2c, **Supplementary Figure 1a**, and **Supplementary Data 1**). The CS distribution was found to be bimodal, with a large fraction of the library centered around a CS value of +13 compared to the non-targeting guide population (CS_norm_). K-means clustering analysis classified the guides into four activity groups based on CS_norm_: highly active (CS_norm_ > 11.46), medium activity (6.90 <CS_norm_< 11.46), low activity (1.36 <CS_norm_< 6.90), and inactive (CS_norm_ < 1.36) guides. Based on this analysis, only 1.3% of the guides in the library are inactive, while 75.6% are highly active (Figure 2d). Active sgRNAs (including low, medium, and high activity) collectively targeted 5,396 genes. Moreover, 83% of the genes were targeted by six active sgRNAs in the library, while 30 genes were not targeted by any active guide. Validation experiments on a subpopulation of guides confirmed that CS is an accurate representation of Cas9 activity (Figure 2e and **Supplementary Figure 2**). With one exception, Cas9-expressing NHEJ-deficient cells expressing twenty-four active sgRNAs exhibited either no or limited growth compared to the empty vector transformation (p < 0.0005). In contrast, 15 of 16 samples with inactive sgRNAs demonstrated growth comparable to the control, thus supporting CS as a quantitative metric for CRISPR-Cas9 activity. Taken together, the CS profiles, library analysis, and CS validation show that the designed library is highly active and has near complete genome-wide coverage of expressed genes.

### 2.3 Activity corrected fitness screens enable accurate essential gene classification

With the CS profile in-hand, we next set out to conduct a fitness screen and determine FS values for every guide in the library and gene in *K. phaffii* GS115 strain (**Supplementary Data 1**). The resulting library FS profile (FS values for every guide) was bimodal with distinctive peaks at FS approximately −2.8 and −8 (Figure 3a). At earlier time points, the FS distribution was less pronounced (**Supplementary Figure 1b**), therefore we used the day-6 time point to define essential genes under glucose growth conditions (2% glucose, SD-H, 30LJ). Using our acCRISPR analysis pipeline (Ramesh et al., 2023), low activity guides were analytically removed from the library before defining FS values per gene and calling essential genes. acCRSIPR identified a CS_threshold_ of 7 to maximize library activity; only guides with a CS value of 7 or greater were used to calculate a gene’s FS value (Figure 3b). At this threshold, the library maintained an average CS value of 8.25, an average of 4.26 guides targeted each gene, and 1604 genes were classified as essential under given growth conditions (corrected p-value < 0.05 per gene against a non-essential gene population, **Supplementary Data 5**). More than 99% of the predicted essential genes were targeted with more than one sgRNA, while genes with only one active sgRNA above the CS_threshold_ were classified as low-confidence essential genes (**Supplementary Figure 3**). In total, 1596 genes were classified as essential with high confidence.

**Figure 3.**
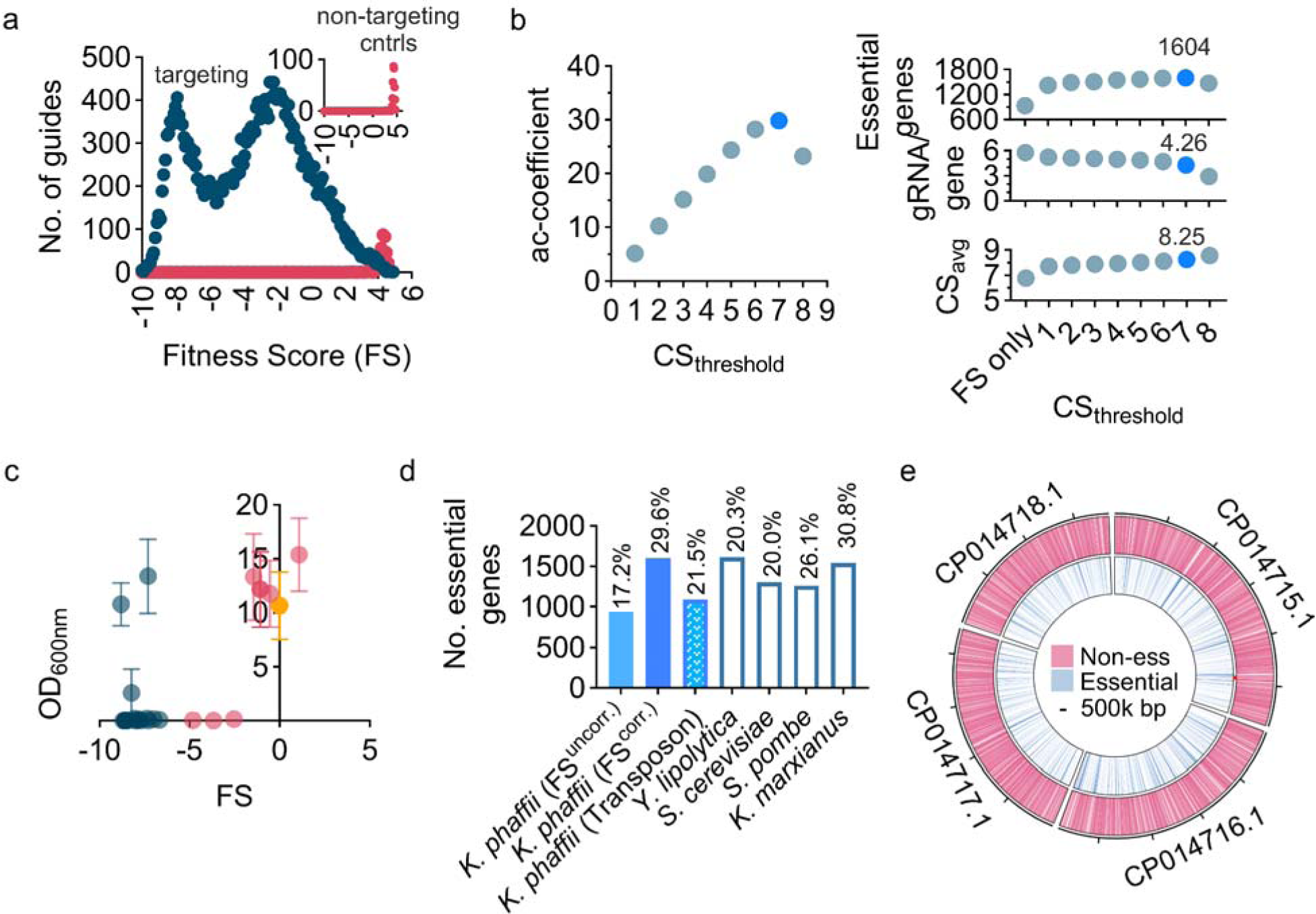
Activity corrected functional genetic screening in *K. phaffii*. **a)** FS frequency distribution per sgRNA. The presented FS values are the mean of three biological replicates per sgRNA at day 6. **b)** Essential gene identification using acCRISPR. The maximum activity correction coefficient (ac-coefficient) occurred at CS_threshold_ value of 7, indicating the conditions for the highest library activity and coverage. At this threshold, 1604 genes were classified as essential (corrected p-value < 0.05). Screens were conducted in SD-H, with 2% glucose, 30 °C. **c)** Individual validation of 17 predicted essential genes (dar blue) and 8 non-essential genes (magenta). A knockout in essential genes leads to low cell viability or cell death compared to a control (yellow). Data points and error bars represent the mean of three biological replicates and one standard deviation three days after subculturing in fresh selective media, respectively. **d)** The number of essential genes in *K. phaffii* with and without activity correction compared with essential gene calls from transposon analysis in *K. phaffii* (Zhu et al., 2018), *Yarrowia lipolytica* (Ramesh et al., 2023), *Saccharomyces cerevisiae (Cherry, 2015)*, *Schizosaccharomyces pombe (Kim et al., 2010),* and *Kluyveromyces marxianus* (**Supplementary Data 6**). Values at the top of each bar represent the percentage of the total number of identified essential genes for each species/method. **e)** Distribution of predicted essential and non-essential genes in GS115’s genome when grown on SD-H media with 2% glucose, 30 °C.

To validate the essentiality of the genes identified by acCRISPR, we selected 17 genes characterized as essential and 8 non-essential genes. Using one highly active guide per gene, we conducted a validation test similar to that conducted for CS validation; guides were transformed into GS115 *his4*::*CAS9* and allowed to grow for up to three days after transferring transformants to fresh selective media. Disruption of an essential gene should produce cultures with no-growth, while disruption of non-essential genes should have minimal effect on culture fitness. Of the 17 essential genes tested, 15 showed no or limited growth compared to the negative control (p < 0.05). Five of eight non-essential gene knockouts grew similar to the negative control (Figure 3c and **Supplementary Figure 4**), while three showed minimal or no growth.

Based on the analysis of model yeast species, roughly 20 to 30% of yeast genes are essential for growth. For example, 19.9% of *S. cerevisiae* genes are classified as essential (Cherry, 2015), while in *S. pombe* an upward of 26.1% of genes are essential (Kim et al., 2010). Our previous analysis of *Yarrowia lipolytica* identified 24.0% of genes as essential for growth on glucose (Ramesh et al., 2023), and a similar analysis of *Kluyveromyces marxianus* suggests that 30.8% of its genes are essential (Figure 3d and **Supplementary Data 6**). Here, we make the comparison to these species as an additional validation step to the essential gene classification in *K. phaffii*. Without activity correction via acCRISPR, only 934 *K. phaffii* genes (17.21% of all CDSs) were identified as essential, suggesting that including all guides in the library results in underestimation of gene essentiality. In addition, a genome-wide transposon insertion library, which is known to under-represent shorter genes (Wang et al., 2018), only identified 1086 essential genes in GS115 with high confidence and an additional 887 with low confidence (Zhu et al., 2018). The activity corrected screens conducted here classified a total of 1604 genes as essential (98.4% high confidence calls) or 29.55% of coding sequences in *K. phaffii* GS115, evenly distributed across the genome (Figure 3e).

We further validated the essential gene set via Gene Ontology (GO) enrichment analysis (**Supplementary Figure 5**) (Ashburner et al., 2000; Harris et al., 2004). The analysis revealed multiple significantly enriched GO terms (adj. p < 0.05; see **Supplementary Data 7** for all GO terms pertaining to molecular function (MF), biological process (BP), cellular component (CC), and KEGG pathway enrichment analysis) with markedly lower FS values compared to the average FS value of all genes. It was anticipated that terms functional for fundamental cell processes would be enriched. As expected genes involved in translation, protein transport and maturation, DNA replication, ribosomal subunit export and assembly, and mitochondrial genes were significantly enriched. Taken together with the other validation methods described above, the essential genes identified from our CRISPR screens represent an accurate classification of essential genes.

### 2.4 Defining a consensus set of essential genes for *Komagataella phaffii* on glucose

The CRISPR-Cas9 screens conducted here, along with the transposon screen conducted by others, provide an opportunity to define a consensus set of essential genes for *K. phaffii* GS115 on glucose. Our validation experiments showed that the activity corrected CRISPR screen yielded a reasonably low false positive rate, but also identified the possibility of false negative (**Supplementary Figure 4**). Given this, we created a consensus set of essential genes by taking the union set called by both technologies (Figure 4a and **Supplementary Data 8**). Among the 1086 high confidence essential genes characterized in the transposon study, 1064 have homologs based on the updated genome annotation used in our study (Alva et al., 2021); only these gene were used to define the consensus set. The union set includes 1880 genes, 816 and 276 of which were only called by the CRISPR screen and the transposon study, respectively, and 788 genes called by both technologies.

**Figure 4.**
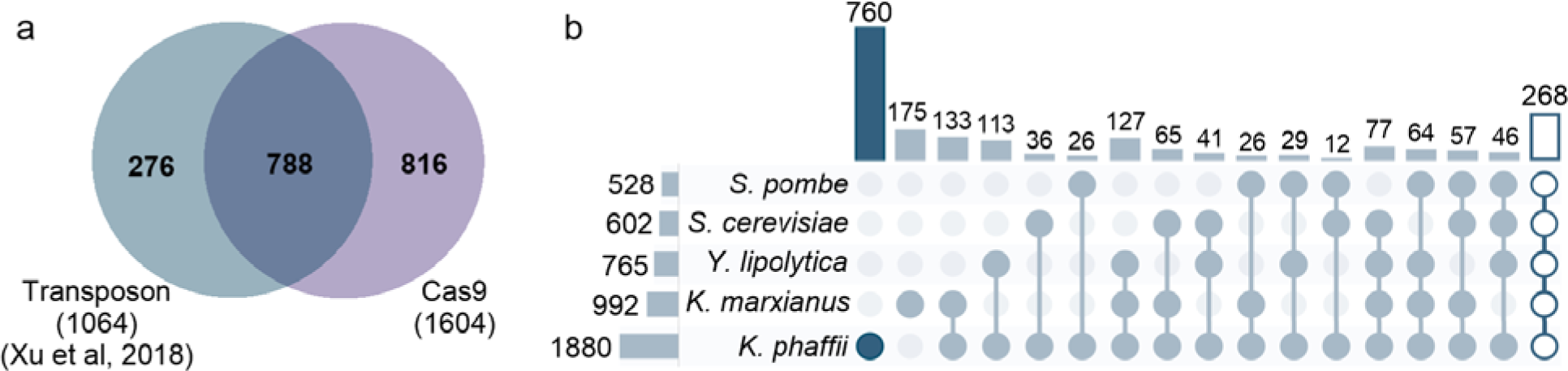
Identification of a consensus set of essential genes for *K. phaffii*. **a)** Venn diagram representation of the number of essential genes identified based on our CRISPR-Cas9 screen, transposon analysis, and their overlap. The consensus essential gene list for *K. phaffii* GS115 on glucose is identified as the union of genes characterized as essential based on CRISPR-Cas9 screens and transposon analysis. **b)** Upset plot representation of the number of essential genes that are common between different yeast species. Values on the top of vertical bars represent the number of essential genes in *K. phaffii* that have essential homologs in other species. Values on the left of the horizontal bars are the intersection of essential genes between species.

The consensus set of 1880 essential genes for *K. phaffii* had 992, 765, 602, and 528 essential homologs in *K. marxianus*, *Y. lipolytica*, *S. cerevisiae* and *S. pombe*, respectively (Figure 4b). Comparison between the consensus set and essential genes in other species also reveals a set of 268 core essential genes common to all five analyzed species as well as 760 genes exclusively essential to *K. phaffii*. According to phylogenetic assessment, the divergence between *S. pombe* and *S. cerevisiae* occurred approximately 420 to 330 million years ago, leading to more genetic distinction among these two species (Sipiczki, 2000). *S. cerevisiae* and *K. phaffii* separated from each other more recently, around 250 million years ago (Bernauer et al., 2020). This relatively more recent divergence likely accounts for the higher number of shared essential genes between *K. phaffii* and *S. cerevisiae* compared to *S. pombe*. *Y. lipolytica*, on the other hand, shares a common ancestor with *K. phaffii (Bernauer et al., 2020)*, potentially contributing to the higher overlap in the number of common essential genes between *K. phaffii* and *Y. lipolytica* compared to the other analyzed species.

### 2.5 Gene Ontology enrichment analysis for essential genes

As additional analysis and validation, Gene Ontology (GO) enrichment and Kyoto Encyclopedia of Genes and Genomes (KEGG) pathway enrichment analyses were conducted for the consensus set of essential genes, the core essential genes common between all five analyzed yeast species, and the essential genes solely belonging to *K. phaffii* (**Supplementary Data 9** and **10)**. Enriched terms (adjusted *p*-value < 0.05) for both analyses, GO terms and KEGG pathways, are represented in Figure 5 and Figure 6.

**Figure 5.**
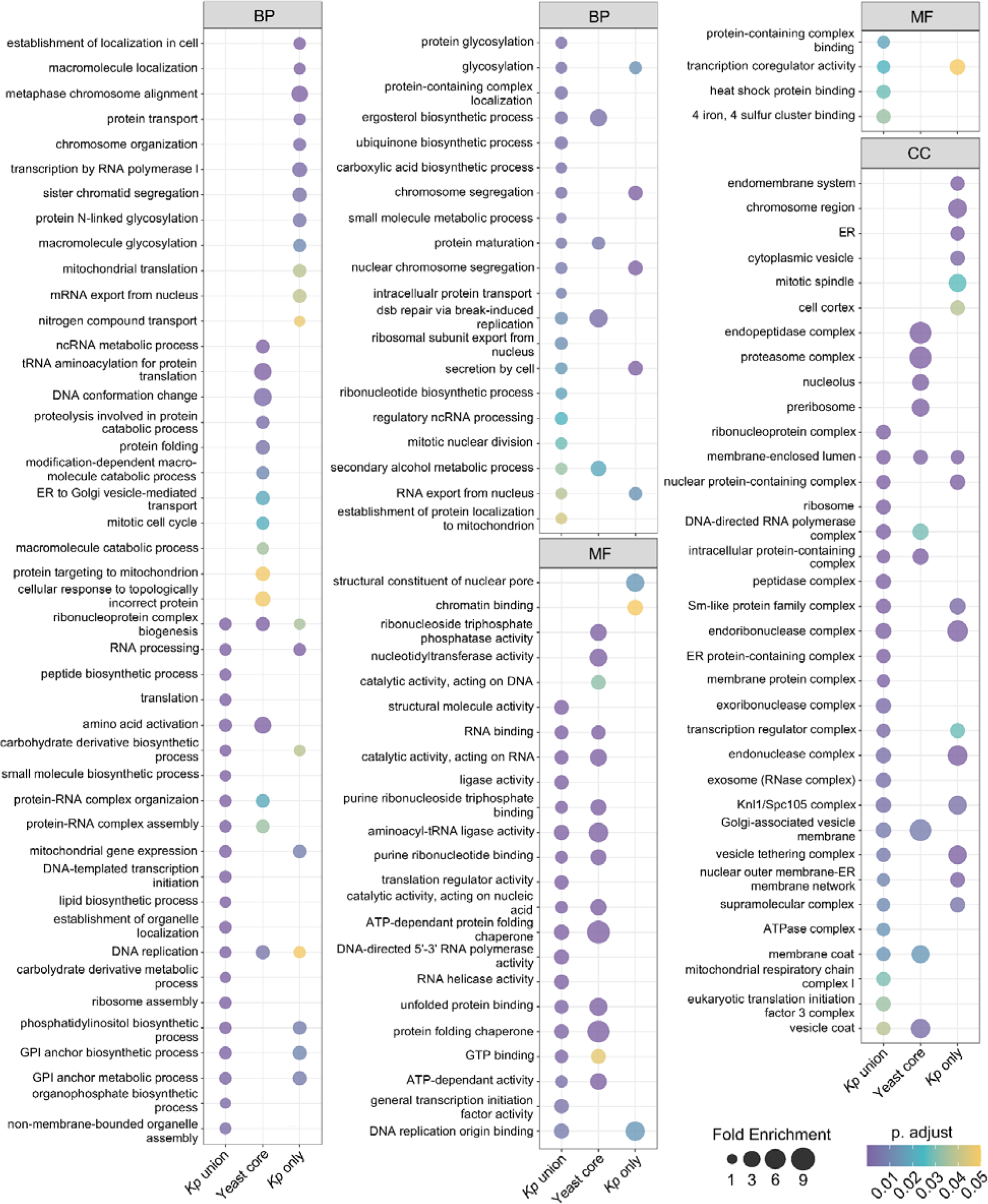
Comparison of functional profiles among different gene sets (Gene Ontology-enrichment analysis) Significantly enriched GO terms in biological process (BP), molecular function (MF), and cellular component (CC) categories (adjusted *p*-value < 0.05) for the consensus set of essential genes (*Kp* union), the core essential genes between five analyzed yeast species (Yeast core), and essential genes solely belong to *K. phaffii* (*Kp* only) are shown. Fold enrichment is the ratio of the frequency of input genes annotated in a term to the frequency of all genes annotated to that term.

**Figure 6.**
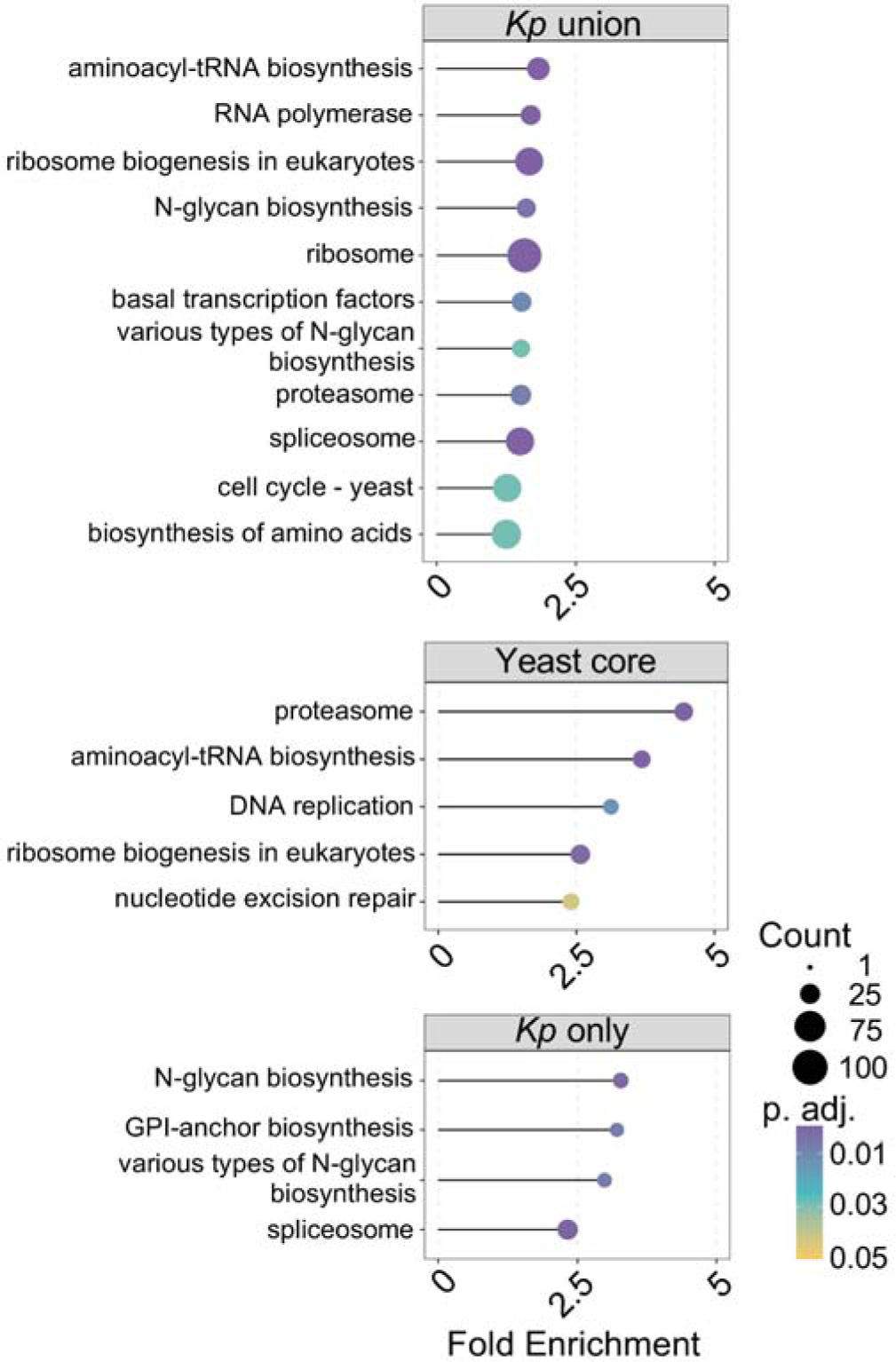
Comparison of functional profiles among different gene sets (Kyoto Encyclopedia of Genes and Genomes pathway analysis) Significantly enriched pathways (adjusted *p*-value < 0.05) for the consensus set of essential genes (*Kp* union), core essential genes between five analyzed species (Yeast core), and essential genes solely belong to *K. phaffii* (*Kp* only). Count represents the number of genes annotated in a specific term, and fold enrichment is defined as the ratio of the frequency of input gene annotated in a term to the frequency of all genes annotated to that term.

As expected, in all three sets of essential genes (consensus, core, and *K. phaffii* specific) we identified vital cell processes and biological pathways (**Supplementary Figure 6-8** and **Supplementary Data 11-13**). The general pattern is that there is minimal overlap of the enriched terms between the three datasets; only three of the 126 enriched GO terms are common between all three sets. These three GO terms are ribonucleoprotein complex biogenesis, DNA replication, and membrane-enclosed lumen. The yeast core enriched terms are most likely composed of conserved genes and pathways preserved through evolution between different species, e.g. tRNA aminoacylation for protein translation, DNA conformation change, protein folding, and cellular response to topologically incorrect proteins. However, terms specifically belonging to *K. phaffii* most likely are composed of genes associated with unique, non-conventional characteristics of the “biotech yeast”.

One of the most important traits of *K. phaffii* is its capability to produce and secrete high titers of recombinant proteins. Various enriched GO terms exclusive to *K. phaffii* in the biological process (BP) category are related to protein production and secretion (Figure 5). Protein transport, establishment of localization in cells, macromolecule localization, and mRNA export from nucleous are amongst the enriched GO terms only for *K. phaffii*. Multiple studies have shown overexpression of genes belonging to these terms to be associated with higher secretion of recombinant products. For instance co-overexpression of *S. cerevisiae* homologs of *SEC63* and *YDJ1* chaperones in *K. phaffii* were attributed to 7.6 times improvement of G-CSF secretion (Zhang et al., 2006). In addition, a study done with a *S. cerevisiae* strain with improved amylase production showed that *ERO1*, *BST1*, *SFB3*, *PEP5, SEC8,* and *EXO84* were upregulated, with all genes being involved in critical roles related to either protein folding or trafficking (Liu et al., 2014). It was suggested that the observed upregulation in these genes might be an indication of the higher activity of the secretory pathway in this strain of *S. cerevisiae*. While these genes were not identified as essential in *S. cerevisiae*, they were categorized as essential in *K. phaffii* based on our screen.

Amongst the GO terms in cellular component (CC) category, endomembrane system, endoplasmic reticulum, and cytoplasmic vesicle are enriched in the *K. phaffii* only data set. Multiple studies have also demonstrated improved protein production with overexpression of genes belonging to these categories. For example, overexpressing the transcription factor *NRG1* in *K. phaffii* is associated with increases in the secretion of Fab2F5, recombinant human trypsinogen, and porcine trypsinogen (Stadlmayr et al., 2010). Another *S. cerevisiae* study, showed that overexpression of the *LHS1* chaperone, which is involved in polypeptide translocation and folding in the ER lumen, increased shake-flask production levels of recombinant human serum albumin, granulocyte-macrophage colony-stimulating factor, and recombinant human transferrin (Payne et al., 2008). Lastly, one study showed the influence of *ERO1* overexpression was able to increase nitrilase production in *K. phaffii* (Shen et al., 2020). Given these examples, the identification of essential genes belonging to specific pathways via genome-wide knockout libraries can be utilized as a novel way to engineer complex phenotypes such as secretion in which overexpression of essential genes can be beneficial.

Another industrially-relevant trait of *K. phaffii* is its ability to glycosylate recombinant proteins (Barone et al., 2023). KEGG pathway enrichment analysis shows N-glycan and various types of N-glycan biosynthesis to be two of the significantly enriched pathways only in *K. phaffii* (Figure 6). Fungi and mammals share initial steps in protein N-glycosylation, including site-specific transfer of a core oligosaccharide (Glc_3_Man_9_GlcNAc_2_) to the nascent polypeptide. Downstream of the first glycosylation events, fungi exhibit a distinct processing pathway in comparison to mammalian cells. Fungi are limited to the addition of mannose and mannosylphosphate sugars to the glycoproteins, which leads to hyper-mannosylation (*S. cerevisiae*) or high-mannose structures (*K. phaffii*) of proteins causing immunogenicity in humans (De Pourcq et al., 2010; Hamilton et al., 2003; Wildt and Gerngross, 2005).

There are 31 genes associated with this pathway in the *K. phaffii* consensus set, among which 17 are exclusively essential in *K. phaffii* including both ER-(*SEC59*, *ALG5*, *ALG13*, *ALG3*, *ALG9*, *ALG12*, *ALG6*, *ALG8*, *OST1*, *OST3*, *SWP1*, *ROT2*, and *DFG10*) and Golgi-residing enzymes (*MNN2*, *MNN11*, MNN10, and *OCH1*). This gene set represents potential important targets for metabolic engineering of non-native glycosylation patterns in *K. phaffii*. For instance, multiple studies have shown that endogenous *OCH1* knockout, a mannosyltransferase which initiates the first step of hypermannosylation in yeast, followed by introducing additional enzymes is crucial in humanizing the glycolysis pathway in *K. phaffii* (Choi et al., 2003; Hamilton et al., 2003; Jacobs et al., 2009). While knocking out *OCH1* would negatively impact cell growth, as indicated by our CRISPR screen and other studies (Dalvie et al., 2022; Moser et al., 2017), the growth impediment is less pronounced in *K. phaffii* compared to *S. cerevisiae*.

Additional distinctive features of *K. phaffii* including the lack of one α-1,3-mannosyltransferase residing in the Golgi, leading to less hyperglycosylation, along with its mammalian-like stacked Golgi structure makes it a superior host for the production of glycoproteins compared to the conventional *S. cerevisiae* system (De Pourcq et al., 2010; Hamilton et al., 2003; Wildt and Gerngross, 2005). Genome-wide knockout libraries thus enable the identification of crucial genes involved in biological pathways, facilitating the understanding of how these pathways differ between microorganisms and offer a novel tool in identification of gene targets to reverse engineer pathways in cells.

## 3.0 Conclusion

High-throughput techniques play a crucial role in advancing metabolic engineering and driving forward genetics. However, these tools are not up to par for non-conventional hosts (Thorwall et al., 2020). Here, we have addressed this issue by designing and validating a redundant 6-fold coverage genome-wide sgRNA library composed of 30,848 active guides that target over 99% of protein coding sequences in the biotech yeast *Komagataella phaffii*. We also optimized the existing transformation protocols for this yeast, enabling the transformation of large-sized libraries for this host. Activity-validated sgRNA libraries can be used to improve screening accuracy, enhance genetic understanding, and aid in optimizing production strains. Notably, similar genome-wide sgRNA libraries have proven effective in finding hits to improve salt tolerance and uncover previously-unknown genes associated with lipid bio-production in *Yarrowia lipolytica* (Ramesh et al., 2023; Schwartz et al., 2019). Application of this tool allowed us to define a consensus set of essential genes for this host on glucose. Through comparison with other known yeast strains, we have identified a set of essential genes exclusive to *K. phaffii*. These essential genes are promising candidates for overexpression, facilitating the engineering of complex phenotypes and advancing metabolic engineering efforts for *K. phaffii*.

## 4.0 Materials and Methods

### 4.1 Strains and culture conditions

*Komagataella phaffii* GS115 (Invitrogen), a strain with histidine deficiency, was used for all experiments (**Supplementary Table 2**). GS115 *his4*::*CAS9* strain was constructed by integrating p*ENO1*-Cas9-PptefT expression cassette into cells’ knocked-out *HIS4* loci. GS115 Δ*KU70* and GS115 *his4*::*CAS9* Δ*KU70* strains were constructed by disrupting *KU70* using CRISPR-Cas9.

All yeast culturing was done at 30 °C in 14 ml polypropylene tubes or in 2 L baffled flasks as noted, at 225 rpm. Under non-selective conditions, yeast strains were initially grown in YPD (1% Bacto yeast extract, 2% Bacto peptone, 2% glucose). Cells were transformed with plasmids expressing sgRNAs and transformants were recovered in histidine-deficient media (SD-his; 0.67% Difco yeast nitrogen base without amino acids, 0.069% CSM-his (Sunrise Science, San Diego, CA), and 2% glucose).

### 4.2 Plasmid construction

All plasmid construction and propagation were conducted in *E. coli* TOP10. Cultures were conducted in Luria-Bertani (LB) broth with either 100LJmgLJ/L ampicillin or 50 mg/L kanamycin at 37 °C in 14 mL polypropylene tubes, at 225 rpm. Plasmids were isolated from *E. coli* cultures using the Zymo Research Plasmid Miniprep Kit.

All plasmids, primers, and sgRNAs used in this work are listed in **Supplementary Tables 3-5**. The D-227 vector containing *CAS9* and a gRNA expression cassette was a kind donation from the Love lab at Massachusetts Institute of Technology (Dalvie et al., 2020). For integration of *CAS9* into *K. phaffii*’s genome, first a highly active *HIS4*-targeting sgRNA was cloned into D-227 vector by digesting the plasmid with BbVCI enzyme (NEB) according to the manufacturer’s instructions. gRNA cloning was carried out according to a protocol developed in the lab previously (Schwartz and Wheeldon, 2018). Primers for sgRNA cloning were obtained from Integrated DNA Technology (IDT). Successful cloning of the sgRNA fragment was confirmed by Sanger sequencing. Next, 1000 bp directly upstream and downstream of the *HIS4*-targeting sgRNA on the genome was PCR amplified and cloned on the upstream and downstream of the p*ENO1*-Cas9-PptefT on D-227 vector using New England BioLabs (NEB) NEBuilder® HiFi DNA Assembly Master Mix. This plasmid was transformed into GS115. *CAS9* integration was verified with PCR amplification of the *HIS4* loci and Sanger sequencing. For all the PCR amplifications in this study Q5 high fidelity polymerase (NEB) was used according to the manufacturer’s instructions.

The backbone of the sgRNA library plasmid (pCRISPRpp) was constructed by PCR amplification of the PARS1 sequence from *K. phaffii*’s genome (Cregg et al., 1985). The E. coli origin of replication and ampicillin resistance gene were PCR amplified from pCRISPRyl (Addgene #70007) (Schwartz et al., 2016). CYC1t and pTEF1 were both PCR amplified from BB3cK_pGAP_23*_pTEF_Cas9 (Addgene #104909) (Gassler et al., 2019). sgRNA expression cassette (ptRNA1_tRNA1_tracrRNA) was PCR amplified from D-227 plasmid. Pp*HIS4* gene was PCR amplified from pMJA089 (Addgene #128518) (Yang et al., 2014). All fragments were cloned to each other to make a single plasmid with NEBuilder® HiFi DNA Assembly Master Mix.

### 4.3 sgRNA library design

CHOPCHOP v3 (Labun et al., 2019) was used to design the sgRNA library for *K. phaffii*. The GS115 reference genome and annotation was downloaded from RefSeq at NCBI (sequence assembly version ASM174695v1, RefSeq assembly accession: GCA_001746955.1) and Bioproject PRJNA669501(Alva et al., 2021). sgRNAs were designed to target the first 300 bp of each coding sequence and tRNA genes to maximize a functional knockout in the gene in case of a CRISPR-induced indel. CHOPCHOP v3 was used to design a preliminary library of 169,034 sgRNAs (**Supplementary Data 3**). Each guide within this preliminary library was characterized by multiple targeting efficiency predictive parameters from various tools including Designer v1 (Doench et al. 2014), Designer v2 (Doench et al. 2016), CRISPRscan (Moreno-Mateos et al. 2015), SSC (Xu et al. 2015), and uCRISPR (Zhang et al., 2019). To enhance the library design process, we also introduced a CS prediction score identified from DeepGuide (Baisya et al. 2022) trained based on *Yarrowia lipolytica* PO1f CRISPR-Cas9 genome-wide sgRNA library CS data. A naive score for each sgRNA was calculated as the aggregate of all the aforementioned normalized targeting efficiency scores.

The uniqueness of each 20 bp sgRNA was analyzed with CHOPCHOP v3 built-in MM0, MM1, MM2, and MM3 scores determining the number of off-target transcripts for each sgRNA with 0, 1, 2, and 3 mismatches, respectively. We also incorporated an extra measure of uniqueness, Seed_MM0, identifying the number of sgRNAs targeting anywhere within the genome with 0 mismatches in the seed region, the last 12 bp of the sgRNA immediately preceding the NGG PAM motif-compared to our sgRNA of interest. Numerous studies have documented that the uniqueness of this seed sequence is a pivotal factor in minimizing the off-target effects of Cas9 (Cong et al., 2013; Hsu et al., 2013; Jiang et al., 2013). Additionally, a self-complementarity score was employed to predict the likelihood of the sgRNA forming a secondary structure with itself, potentially reducing the targeting efficiency (Labun et al., 2019). A comprehensive quality score was assigned to each sgRNA taking into account all uniqueness and self-complementarity scores. A quality score of 1 signifies an sgRNA that not only possesses uniqueness in both its 20 base pair sequence and seed region but also exhibits a minimal likelihood of forming secondary structures. The detailed breakdown of all the defined quality scores can be found in **Supplementary Table 6**.

sgRNAs designed for each coding sequence were initially ranked based on their “quality” score and then the top sgRNAs with the highest “naive” score were chosen for the final library for each coding sequence or tRNA gene. Over 99% of the sgRNAs in the library had a quality score of 1 (**Supplementary Table 7**). Three hundred and fifty sgRNAs with random sequences were also included as non-targeting controls (**Supplementary Data 4**). All designed sgRNAs along with additional data are available in **Supplementary Data 2.**

### 4.4 sgRNA library cloning

60mer linkers were added 5’ and 3’ of each designed sgRNA enabling assembly into pCRISPRpp (**Supplementary Table 4**) and obtained as a pooled oligonucleotide library (Twist BioScience, CA, USA). The library was amplified for 9 cycles with Kapa polymerase (Roche) using the oligonucleotides 5’ tagtggtagaaccaccgcttgtc and 5’ actttttcaagttgataacggactagcc and assembled into pCRISPRpp linearized by BbvCI digestion, and dephosphorylated with quick CIP, using the NEBuilder Hi-Fi Assembly kit (New England Biolabs). To ensure representation of all variants in the population, >330,000 colonies were obtained (*i.e.*, >10 colonies per sgRNA), and the library validated by insert PCR amplification with the oligonucleotides 5’ agccaatcctactacattgatccg and 5’ gtcatgataataatggtttcttagacg. The amplicon library was sequenced on the Illumina MiSeq platform and the data analyzed using custom library quality control pipelines (**Supplementary Data 14**).

### 4.5 Yeast transformation and screening

Transformation of *K. phaffii* was done using a previously described method, with slight modifications (Wu and Letchworth, 2004). Two mL of YPD was inoculated with a single colony of the strain of interest and grown in a 14 mL tube with shaking at 225 rpm overnight. 4 × 10^7^ cells were transferred to 150 ml of YPD in a 500 ml baffled shake flask and grown for ∼14 h (until the culture reached a final OD_600_ = 1.8). 100 ml of cells were chilled on ice for 1.5 h, washed with 1 M ice-cold sorbitol three times, incubated with 25 ml of pretreatment solution (0.1 M lithium acetate (LiAc), 30 mM dithiothreitol (DTT), 0.6 M sorbitol, and 10 mM tris-HCl pH=7.5) for 30 minutes at room temperature, and washed three more times with 1 M ice-cold sorbitol. For each transformation, 8 × 10^8^ cells were mixed with 1 µg of library to a final volume of 80 μl, incubated on ice for 15 min, and pulsed at 1.5 kV with Bio-Rad MicroPulser Electroporator in an ice-cold 0.2-cm-gap cuvette. Immediately after electroporation shock, 1 ml ice-cold solution of YPD and 1 M sorbitol was added to each cuvette. Cells were transferred to 1 ml YPD and 1 M sorbitol in 14 ml tubes, incubated for 3 h at 30°C and 225 rpm for recovery, washed with 1 ml of room-temperature autoclaved water to get rid of the excess plasmid DNA in samples, and transferred to selective media. All centrifugation was done at 3000×g for 5 minutes at 4 LJ. The relationship between cell number and OD_600_ was calculated according to 1 OD_600_ = 5 × 10^7^ cells/mL.

For library transformations, 10 separate transformations were pooled together after recovery to maintain library representation (100-fold coverage, total transformants per biological replicate). Pooled transformants were transferred to 750 ml SD-his for outgrowth experiments in a 2 L baffled shake flask. Three biological replicates were performed for each strain. Transformation efficiency for each replicate and strain is presented in **Supplementary Table 8**. Cells reached to confluency after 3 days (OD_600_ ≈ 8). 1 ml of cells were transferred to 50 ml fresh SD-his in 250 ml baffled shake flasks to perform outgrowth experiments and were allowed to grow for three more days. The experiment was stopped after reaching confluency again on day six of the screen. At each time point, 1 ml of culture was stored at −80 °C to isolate sgRNA expression plasmids for deep sequencing.

### 4.6 Library isolation and sequencing

Frozen culture samples from pooled screens were thawed. Plasmids were isolated from each sample using a Zymo Yeast Plasmid Miniprep Kit (Zymo Research). 500 µL of each sample was divided into two tubes to account for the capacity of the yeast miniprep kit, specifically to ensure complete lysis of the cells using Zymolyase. The split miniprepped samples from a single strain and replicate were pooled again, and the plasmid copy number was quantified using quantitative PCR with qPCR_GW.F, qPCR_GW.R, and SsoAdvanced Universal SYBR Green Supermix (Bio-Rad). Each pooled sample was confirmed to contain at least 10^7^ plasmids ensuring sufficient coverage of the sgRNA library. Recovered plasmid copy number and coverage for each sample and replicate is presented in **Supplementary Table 9**.

To prepare samples for next generation sequencing (NGS), isolated plasmids from each sample were used as PCR templates using forward (NGS1-4.F) and reverse primers (NGS1-9.R). Different forward and reverse barcodes and pseudo-barcodes were used in primers to increase complexity for NGS and to enable us to differentiate between samples later on. NGS primers were ordered as Ultramer DNA oligos from IDT. At least 0.5 ng of the recovered plasmids (∼ molecules) were used to amplify the amplicons in a 16-cycle PCR reaction to minimize any bias. PCR products were cleaned by a double-sided cleanup technique using AMPure XP beads and tested with a Bioanalyzer to ensure the correct length of amplicons. 80 nmol of FS and cutting score CS samples were pooled together separately and submitted for sequencing on a NextSeq2000 using a P2 100 cycle kit.

### 4.7 Generating sgRNA read counts from raw reads

Next generation sequencing files were processed with custom python codes. Read quality was analyzed using FastQC v0.11.9. Raw reads were demultiplexed and truncated using Cutadapt 4.2 (Martin, 2011) to only include the sgRNAs. sgRNA abundance in sequencing reads were initially calculated using naïve exact matching (NEM). Inexact matching (IEM) via Bowtie v1.3.1 was performed for those reads that were not aligned to the library with NEM. A large proportion of the counts were calculated from exact matching. A total of 394 sgRNAs had zero counts or had very low normalized abundance (<1% of the normalized mean abundance of the library) based on NGS data obtained from extracted plasmids from cells. Therefore, these sgRNAs were removed from further analysis. Normalized sgRNA read count between biological replicates was plotted to verify consistency and reproducibility of the experiments (**Supplementary Table 10**).

### 4.8 Cutting and Fitness score calculations

Based on our acCRISPR analysis pipeline (Ramesh et al., 2023), fitness score (FS) and cutting score (CS) was calculated by first adding a pseudo-count of one to each raw count before normalization. The read counts for each sgRNA were normalized to the total number of reads for that specific sample. Fitness score value for each sgRNA was calculated as the log_2_ ratio of normalized read counts obtained in GS115 *his4*::*CAS9* to normalized counts in GS115 strain. Similarly, cutting score (CS) was defined as the −log_2_ ratio of normalized reads obtained in GS115 *his4*::*CAS9* Δ*KU70* to counts in GS115 Δ*KU70* (**Supplementary Data 1**).

### 4.9 Essential gene identification

FS and CS values of sgRNA at day six were used as input to acCRISPR v1.0.0 (Ramesh et al., 2023) to identify essential genes from the screen. A CS_threshold_ of 7.0 was used to remove low-activity sgRNAs from the original library, due to the maximum value of ac-coefficient at this threshold. FS of a gene was computed by acCRISPR as the average of FS of all sgRNAs with CS above 7.0 targeting that gene. Genes having FDR-corrected p < 0.05 were deemed as essential.

### 4.10 Finding essential gene homologs in S. cerevisiae, S. pombe, Y. lipolytica, and K. marxianus

Sequences of genes essential identified in this study and/or in the transposon study (Zhu et al., 2018) were aligned to genes in *S. cerevisiae*, *S. pombe,* and *Y. lipolytica* using BLASTP, and to genes in *K. marxianus* using TBLASTN. *S. cerevisiae* essential genes (phenotype:inviable) were retrieved from the *Saccharomyces* Genome Database (SGD), *S. pombe* essential genes were taken from (Kim et al., 2010), and *Y. lipolytica* essential genes were taken from the consensus set defined in (Ramesh et al., 2023). *K. marxianus* essential genes were identified from a CRISPR-Cas9 genome-wide library in the CBS6556 strain in our lab. Pairs of query and subject sequences having > 40% identity from BLAST were deemed as homologs.

### 4.11 Experimental validation of fitness and cutting scores

Selected genes/sgRNAs were chosen for essential gene/cutting score validations, respectively. The essential gene validations were done by performing a single-gene knockout using high cutting score sgRNAs targeting 17 predicted essential genes and 8 non-essential genes in the GS115 *his4*::*CAS9* background. Additionally, sixteen inactive (CS_norm_ < 1.36), four low-activity (1.36 <CS_norm_< 6.90), two medium-activity (6.90 <CS_norm_< 11.46), and sixteen high-activity (CS_norm_ > 11.46) sgRNAs were chosen for CS validations in the GS115 *his4*::*CAS9 ku70* background. Individual plasmids containing sgRNAs were cloned as was mentioned previously. Transformants were grown in 4 mL of SD-his for two days, followed by sub-culturing in 2 mL of fresh selective media. The OD_600_ of the samples were measured three days after the sub-culture. All of the validation experiments were done in three biological replicates.

### 4.12 Functional enrichment analysis

The organism package for *K. phaffii* GS115 (NCBI Taxonomy ID : 644223) was created using the AnnotationForge package (version 1.44.0) (Marc Carlson, 2017). ClusterProfiler package (version 4.10.0) was used for functional enrichment analysis (Wu et al., 2021; Yu et al., 2012). GO (Gene Ontology) and KEGG (Kyoto Encyclopedia of Genes and Genomes) pathway enrichment analysis were applied to the genes in the consensus set, yeast core set, and *K. phaffii*-specific set. The *p*-value was calculated by Fisher’s exact test. Benjamini-Hochberg procedure was applied to correct *p*-values. Significant GO terms and pathways were identified with a cutoff for adjusted *p*-value (adj. *p*-value < 0.05). Fold enrichments, defined as the ratio of the frequency of input genes annotated in a term to the frequency of all genes annotated to that term, for all the enriched terms were also calculated to interpret the results better. To get a more effective interpretation from the analysis, some redundant GO terms (with semantic similarities over 0.7) were removed by applying the *simplify* function in the ClusterProfiler package (version 4.10.0). For some GO terms with a parent-child semantic relationship having the same *p*-values and geneRatio (ratio of input genes annotated in a term), the parent terms were eliminated from the list.

## Author contributions

AT, VT, and IW conceived the project, planned the experiments, and analyzed the data. *s*, respectively. VT, AT, ML, and ZX analyzed the essential genes. ZX and SC did the ClusterProfiler analysis. IB and RM created the sgRNA library. AT, VT, ZX, IB and IW wrote the manuscript with input from all authors.

## Acknowledgments

This work was supported by NSF-1951942, NSF-2225878, and NSF-1922642. The work (proposal: https://doi.org/10.46936/10.25585/60001310) conducted by the U.S. Department of Energy Joint Genome Institute (https://ror.org/04xm1d337), a DOE Office of Science User Facility, is supported by the Office of Science of the U.S. Department of Energy operated under Contract No. DE-AC02-05CH11231. We thank Dipankar Baisya for his assistance with DeepGuide and Adithya Ramesh for guidance with the library design. We also thank Dr. J. Christopher Love for providing the Cas9 expressing plasmid, D-227.

## Data availability

The demultiplexed sgRNA sequencing data for our CRISPR-Cas9 screens generated for this study have been deposited in the NCBI SRA database under accession code PRJNA1068757. Source data for main figures in the study not included in **Supplementary Data 1-14** is provided in **Supplementary Data 15**. Any remaining information can be obtained from the corresponding author upon reasonable request.

## Code availability

Source code for the CRISPR-Cas9 library design can be found at https://github.com/ianwheeldon/Kphaffii_library_design.git/. Custom python scripts that were used for the processing of Illumina reads to generate sgRNA abundance for the Cas9 screens can also be found at the same link.

## References

Ahmad, M., Hirz, M., Pichler, H., Schwab, H., 2014. Protein expression in Pichia pastoris: recent achievements and perspectives for heterologous protein production. Appl. Microbiol. Biotechnol. 98, 5301–5317.

Alva, T.R., Riera, M., Chartron, J.W., 2021. Translational landscape and protein biogenesis demands of the early secretory pathway in Komagataella phaffii. Microb. Cell Fact. 20, 19.

Ashburner, M., Ball, C.A., Blake, J.A., Botstein, D., Butler, H., Cherry, J.M., Davis, A.P., Dolinski, K., Dwight, S.S., Eppig, J.T., Harris, M.A., Hill, D.P., Issel-Tarver, L., Kasarskis, A., Lewis, S., Matese, J.C., Richardson, J.E., Ringwald, M., Rubin, G.M., Sherlock, G., 2000. Gene ontology: tool for the unification of biology. The Gene Ontology Consortium. Nat. Genet. 25, 25–29.

Ata, Ö., Ergün, B.G., Fickers, P., Heistinger, L., Mattanovich, D., Rebnegger, C., Gasser, B., 2021. What makes Komagataella phaffii non-conventional? FEMS Yeast Res. 21. 10.1093/femsyr/foab059

Baisya, D., Ramesh, A., Schwartz, C., Lonardi, S., Wheeldon, I., 2022. Genome-wide functional screens enable the prediction of high activity CRISPR-Cas9 and -Cas12a guides in Yarrowia lipolytica. Nat. Commun. 13, 922.

Barone, G.D., Emmerstorfer-Augustin, A., Biundo, A., Pisano, I., Coccetti, P., Mapelli, V., Camattari, A., 2023. Industrial Production of Proteins with Pichia pastoris-Komagataella phaffii. Biomolecules 13. 10.3390/biom13030441

Bernauer, L., Radkohl, A., Lehmayer, L.G.K., Emmerstorfer-Augustin, A., 2020. Komagataella phaffii as Emerging Model Organism in Fundamental Research. Front. Microbiol. 11, 607028.

Cai, P., Duan, X., Wu, X., Gao, L., Ye, M., Zhou, Y.J., 2021. Recombination machinery engineering facilitates metabolic engineering of the industrial yeast Pichia pastoris. Nucleic Acids Res. 49, 7791–7805.

Cherry, J.M., 2015. The Saccharomyces Genome Database: Advanced Searching Methods and Data Mining. Cold Spring Harb. Protoc. 2015, db.prot088906.

Choi, B.-K., Bobrowicz, P., Davidson, R.C., Hamilton, S.R., Kung, D.H., Li, H., Miele, R.G., Nett, J.H., Wildt, S., Gerngross, T.U., 2003. Use of combinatorial genetic libraries to humanize N-linked glycosylation in the yeast *Pichia pastoris*. Proc. Natl. Acad. Sci. U. S. A. 100, 5022– 5027.

Cong, L., Ran, F.A., Cox, D., Lin, S., Barretto, R., Habib, N., Hsu, P.D., Wu, X., Jiang, W., Marraffini, L.A., Zhang, F., 2013. Multiplex genome engineering using CRISPR/Cas systems. Science 339, 819–823.

Cregg, J.M., Barringer, K.J., Hessler, A.Y., Madden, K.R., 1985. Pichia pastoris as a host system for transformations. Mol. Cell. Biol. 5, 3376–3385.

Dalvie, N.C., Leal, J., Whittaker, C.A., Yang, Y., Brady, J.R., Love, K.R., Love, J.C., 2020. Host-Informed Expression of CRISPR Guide RNA for Genomic Engineering in Komagataella phaffii. ACS Synth. Biol. 9, 26–35.

Dalvie, N.C., Lorgeree, T., Biedermann, A.M., Love, K.R., Love, J.C., 2022. Simplified Gene Knockout by CRISPR-Cas9-Induced Homologous Recombination. ACS Synth. Biol. 11, 497–501.

Daly, R., Hearn, M.T.W., 2005. Expression of heterologous proteins in Pichia pastoris: a useful experimental tool in protein engineering and production. J. Mol. Recognit. 18, 119–138.

De Pourcq, K., De Schutter, K., Callewaert, N., 2010. Engineering of glycosylation in yeast and other fungi: current state and perspectives. Appl. Microbiol. Biotechnol. 87, 1617–1631.

Dong, C., Fu, S., Karvas, R.M., Chew, B., Fischer, L.A., Xing, X., Harrison, J.K., Popli, P., Kommagani, R., Wang, T., Zhang, B., Theunissen, T.W., 2022. A genome-wide CRISPR-Cas9 knockout screen identifies essential and growth-restricting genes in human trophoblast stem cells. Nat. Commun. 13, 2548.

Gasser, B., Prielhofer, R., Marx, H., Maurer, M., Nocon, J., Steiger, M., Puxbaum, V., Sauer, M., Mattanovich, D., 2013. Pichia pastoris: protein production host and model organism for biomedical research. Future Microbiol. 8, 191–208.

Gassler, T., Heistinger, L., Mattanovich, D., Gasser, B., Prielhofer, R., 2019. CRISPR/Cas9-Mediated Homology-Directed Genome Editing in Pichia pastoris. Methods Mol. Biol. 1923, 211–225.

Guo, Y., Park, J.M., Cui, B., Humes, E., Gangadharan, S., Hung, S., FitzGerald, P.C., Hoe, K.-L., Grewal, S.I.S., Craig, N.L., Levin, H.L., 2013. Integration profiling of gene function with dense maps of transposon integration. Genetics 195, 599–609.

Hamilton, S.R., Bobrowicz, P., Bobrowicz, B., Davidson, R.C., Li, H., Mitchell, T., Nett, J.H., Rausch, S., Stadheim, T.A., Wischnewski, H., Wildt, S., Gerngross, T.U., 2003. Production of complex human glycoproteins in yeast. Science 301, 1244–1246.

Harris, M.A., Clark, J., Ireland, A., Lomax, J., Ashburner, M., Foulger, R., Eilbeck, K., Lewis, S., Marshall, B., Mungall, C., Richter, J., Rubin, G.M., Blake, J.A., Bult, C., Dolan, M., Drabkin, H., Eppig, J.T., Hill, D.P., Ni, L., Ringwald, M., Balakrishnan, R., Cherry, J.M., Christie, K.R., Costanzo, M.C., Dwight, S.S., Engel, S., Fisk, D.G., Hirschman, J.E., Hong, E.L., Nash, R.S., Sethuraman, A., Theesfeld, C.L., Botstein, D., Dolinski, K., Feierbach, B., Berardini, T., Mundodi, S., Rhee, S.Y., Apweiler, R., Barrell, D., Camon, E., Dimmer, E., Lee, V., Chisholm, R., Gaudet, P., Kibbe, W., Kishore, R., Schwarz, E.M., Sternberg, P., Gwinn, M., Hannick, L., Wortman, J., Berriman, M., Wood, V., de la Cruz, N., Tonellato, P., Jaiswal, P., Seigfried, T., White, R., Gene Ontology Consortium, 2004. The Gene Ontology (GO) database and informatics resource. Nucleic Acids Res. 32, D258–61.

Hsu, P.D., Scott, D.A., Weinstein, J.A., Ran, F.A., Konermann, S., Agarwala, V., Li, Y., Fine, E.J., Wu, X., Shalem, O., Cradick, T.J., Marraffini, L.A., Bao, G., Zhang, F., 2013. DNA targeting specificity of RNA-guided Cas9 nucleases. Nat. Biotechnol. 31, 827–832.

Jacobs, P.P., Geysens, S., Vervecken, W., Contreras, R., Callewaert, N., 2009. Engineering complex-type N-glycosylation in Pichia pastoris using GlycoSwitch technology. Nat. Protoc. 4, 58–70.

Jiang, W., Bikard, D., Cox, D., Zhang, F., Marraffini, L.A., 2013. RNA-guided editing of bacterial genomes using CRISPR-Cas systems. Nat. Biotechnol. 31, 233–239.

Jinek, M., Chylinski, K., Fonfara, I., Hauer, M., Doudna, J.A., Charpentier, E., 2012. A Programmable Dual-RNA–Guided DNA Endonuclease in Adaptive Bacterial Immunity. Science 337, 816–821.

Kim, D.-U., Hayles, J., Kim, D., Wood, V., Park, H.-O., Won, M., Yoo, H.-S., Duhig, T., Nam, M., Palmer, G., Han, S., Jeffery, L., Baek, S.-T., Lee, H., Shim, Y.S., Lee, M., Kim, L., Heo, K.-S., Noh, E.J., Lee, A.-R., Jang, Y.-J., Chung, K.-S., Choi, S.-J., Park, J.-Y., Park, Y., Kim, H.M., Park, S.-K., Park, H.-J., Kang, E.-J., Kim, H.B., Kang, H.-S., Park, H.-M., Kim, K., Song, K., Song, K.B., Nurse, P., Hoe, K.-L., 2010. Analysis of a genome-wide set of gene deletions in the fission yeast Schizosaccharomyces pombe. Nat. Biotechnol. 28, 617–623.

Labun, K., Montague, T.G., Krause, M., Torres Cleuren, Y.N., Tjeldnes, H., Valen, E., 2019. CHOPCHOP v3: expanding the CRISPR web toolbox beyond genome editing. Nucleic Acids Res. 47, W171–W174.

Liu, R., Chen, L., Jiang, Y., Zhou, Z., Zou, G., 2015. Efficient genome editing in filamentous fungus Trichoderma reesei using the CRISPR/Cas9 system. Cell Discov 1, 15007.

Liu, Z., Liu, L., Österlund, T., Hou, J., Huang, M., Fagerberg, L., Petranovic, D., Uhlén, M., Nielsen, J., 2014. Improved production of a heterologous amylase in Saccharomyces cerevisiae by inverse metabolic engineering. Appl. Environ. Microbiol. 80, 5542–5550.

Löbs, A.-K., Engel, R., Schwartz, C., Flores, A., Wheeldon, I., 2017a. CRISPR-Cas9-enabled genetic disruptions for understanding ethanol and ethyl acetate biosynthesis in Kluyveromyces marxianus. Biotechnol. Biofuels 10, 164.

Löbs, A.-K., Schwartz, C., Wheeldon, I., 2017b. Genome and metabolic engineering in non-conventional yeasts: Current advances and applications. Synth Syst Biotechnol 2, 198– 207.

Love, K.R., Dalvie, N.C., Love, J.C., 2018. The yeast stands alone: the future of protein biologic production. Curr. Opin. Biotechnol. 53, 50–58.

Lupish, B., Hall, J., Schwartz, C., Ramesh, A., Morrison, C., Wheeldon, I., 2022. Genome-wide CRISPR-Cas9 screen reveals a persistent null-hyphal phenotype that maintains high carotenoid production in Yarrowia lipolytica. Biotechnol. Bioeng. 119, 3623–3631.

Marc Carlson, H.P., 2017. AnnotationForge. Bioconductor. 10.18129/B9.BIOC.ANNOTATIONFORGE

Martin, M., 2011. Cutadapt removes adapter sequences from high-throughput sequencing reads. EMBnet.journal 17, 10–12.

Michel, A.H., Hatakeyama, R., Kimmig, P., Arter, M., Peter, M., Matos, J., De Virgilio, C., Kornmann, B., 2017. Functional mapping of yeast genomes by saturated transposition. Elife 6. 10.7554/eLife.23570

Moreb, E.A., Lynch, M.D., 2021. Genome dependent Cas9/gRNA search time underlies sequence dependent gRNA activity. Nat. Commun. 12, 5034.

Morgens, D.W., Deans, R.M., Li, A., Bassik, M.C., 2016. Systematic comparison of CRISPR/Cas9 and RNAi screens for essential genes. Nat. Biotechnol. 34, 634–636.

Moser, J.W., Wilson, I.B.H., Dragosits, M., 2017. The adaptive landscape of wildtype and glycosylation-deficient populations of the industrial yeast Pichia pastoris. BMC Genomics 18, 597.

Patterson, K., Yu, J., Landberg, J., Chang, I., Shavarebi, F., Bilanchone, V., Sandmeyer, S., 2018. Functional genomics for the oleaginous yeast Yarrowia lipolytica. Metab. Eng. 48, 184–196.

Payne, T., Finnis, C., Evans, L.R., Mead, D.J., Avery, S.V., Archer, D.B., Sleep, D., 2008. Modulation of chaperone gene expression in mutagenized Saccharomyces cerevisiae strains developed for recombinant human albumin production results in increased production of multiple heterologous proteins. Appl. Environ. Microbiol. 74, 7759–7766.

Ramesh, A., Trivedi, V., Lee, S., Tafrishi, A., Schwartz, C., Mohseni, A., Li, M., Lonardi, S., Wheeldon, I., 2023. acCRISPR: an activity-correction method for improving the accuracy of CRISPR screens. Commun Biol 6, 617.

Schwartz, C., Cheng, J.-F., Evans, R., Schwartz, C.A., Wagner, J.M., Anglin, S., Beitz, A., Pan, W., Lonardi, S., Blenner, M., Alper, H.S., Yoshikuni, Y., Wheeldon, I., 2019. Validating genome-wide CRISPR-Cas9 function improves screening in the oleaginous yeast Yarrowia lipolytica. Metab. Eng. 55, 102–110.

Schwartz, C.M., Hussain, M.S., Blenner, M., Wheeldon, I., 2016. Synthetic RNA Polymerase III Promoters Facilitate High-Efficiency CRISPR-Cas9-Mediated Genome Editing in Yarrowia lipolytica. ACS Synth. Biol. 5, 356–359.

Schwartz, C., Wheeldon, I., 2018. CRISPR-Cas9-Mediated Genome Editing and Transcriptional Control in Yarrowia lipolytica. Methods Mol. Biol. 1772, 327–345.

Shalem, O., Sanjana, N.E., Hartenian, E., Shi, X., Scott, D.A., Mikkelson, T., Heckl, D., Ebert, B.L., Root, D.E., Doench, J.G., Zhang, F., 2014. Genome-scale CRISPR-Cas9 knockout screening in human cells. Science 343, 84–87.

Shen, Q., Yu, Z., Lv, P.-J., Li, Q., Zou, S.-P., Xiong, N., Liu, Z.-Q., Xue, Y.-P., Zheng, Y.-G., 2020. Engineering a Pichia pastoris nitrilase whole cell catalyst through the increased nitrilase gene copy number and co-expressing of ER oxidoreductin 1. Appl. Microbiol. Biotechnol. 104, 2489–2500.

Sipiczki, M., 2000. Where does fission yeast sit on the tree of life? Genome Biol. 1, REVIEWS1011.

Stadlmayr, G., Benakovitsch, K., Gasser, B., Mattanovich, D., Sauer, M., 2010. Genome-scale analysis of library sorting (GALibSo): Isolation of secretion enhancing factors for recombinant protein production in Pichia pastoris. Biotechnol. Bioeng. 105, 543–555.

Thorwall, S., Schwartz, C., Chartron, J.W., Wheeldon, I., 2020. Stress-tolerant non-conventional microbes enable next-generation chemical biosynthesis. Nat. Chem. Biol. 16, 113–121.

Trivedi, V., Ramesh, A., Wheeldon, I., 2023. Analyzing CRISPR screens in non-conventional microbes. J. Ind. Microbiol. Biotechnol. 50. 10.1093/jimb/kuad006

Wang, T., Guan, C., Guo, J., Liu, B., Wu, Y., Xie, Z., Zhang, C., Xing, X.-H., 2018. Pooled CRISPR interference screening enables genome-scale functional genomics study in bacteria with superior performance. Nat. Commun. 9, 2475.

Weninger, A., Hatzl, A.-M., Schmid, C., Vogl, T., Glieder, A., 2016. Combinatorial optimization of CRISPR/Cas9 expression enables precision genome engineering in the methylotrophic yeast Pichia pastoris. J. Biotechnol. 235, 139–149.

Wildt, S., Gerngross, T.U., 2005. The humanization of N-glycosylation pathways in yeast. Nat. Rev. Microbiol. 3, 119–128.

Wu, S., Letchworth, G.J., 2004. High efficiency transformation by electroporation of Pichia pastoris pretreated with lithium acetate and dithiothreitol. Biotechniques 36, 152–154.

Wu, T., Hu, E., Xu, S., Chen, M., Guo, P., Dai, Z., Feng, T., Zhou, L., Tang, W., Zhan, L., Fu, X., Liu, S., Bo, X., Yu, G., 2021. clusterProfiler 4.0: A universal enrichment tool for interpreting omics data. Innovation (Camb) 2, 100141.

Yang, J., Nie, L., Chen, B., Liu, Y., Kong, Y., Wang, H., Diao, L., 2014. Hygromycin-resistance vectors for gene expression in Pichia pastoris. Yeast 31, 115–125.

Yu, G., Wang, L.-G., Han, Y., He, Q.-Y., 2012. clusterProfiler: an R package for comparing biological themes among gene clusters. OMICS 16, 284–287.

Zhang, G., Luo, Y., Dai, X., Dai, Z., 2023. Benchmarking deep learning methods for predicting CRISPR/Cas9 sgRNA on- and off-target activities. Brief. Bioinform. 24. 10.1093/bib/bbad333

Zhang, W., Zhao, H.-L., Xue, C., Xiong, X.-H., Yao, X.-Q., Li, X.-Y., Chen, H.-P., Liu, Z.-M., 2006. Enhanced secretion of heterologous proteins in Pichia pastoris following overexpression of Saccharomyces cerevisiae chaperone proteins. Biotechnol. Prog. 22, 1090–1095.

Zhu, J., Gong, R., Zhu, Q., He, Q., Xu, N., Xu, Y., Cai, M., Zhou, X., Zhang, Y., Zhou, M., 2018. Genome-Wide Determination of Gene Essentiality by Transposon Insertion Sequencing in Yeast Pichia pastoris. Sci. Rep. 8, 10223.

