## Supplementary Material for "Functional genomic screening in *Komagataella phaffii* enabled by high-activity CRISPR-Cas9 library"

### Sanofi, San Diego, CA USA

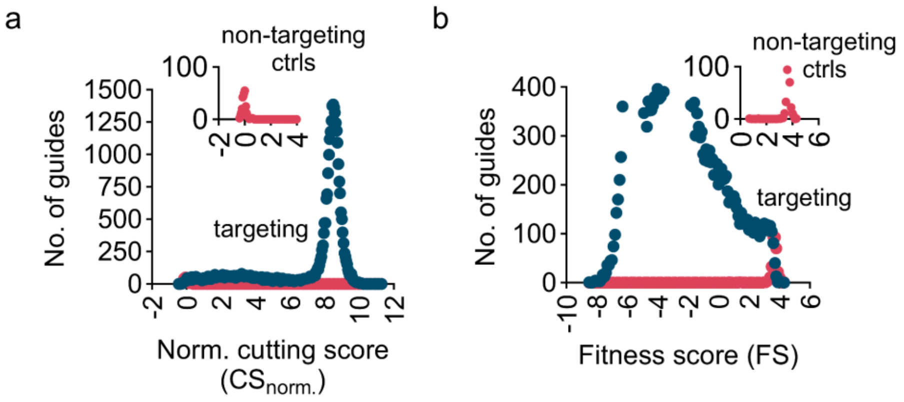


**Supplementary Figure 1** - Frequency distribution of **a)** normalized cutting score (CS); and **b)** fitness score (FS) data at day 3. The presented CS and FS values are the mean of three biological replicates. The presented CS for each sgRNA is normalized to the average CS of non-targeting controls.

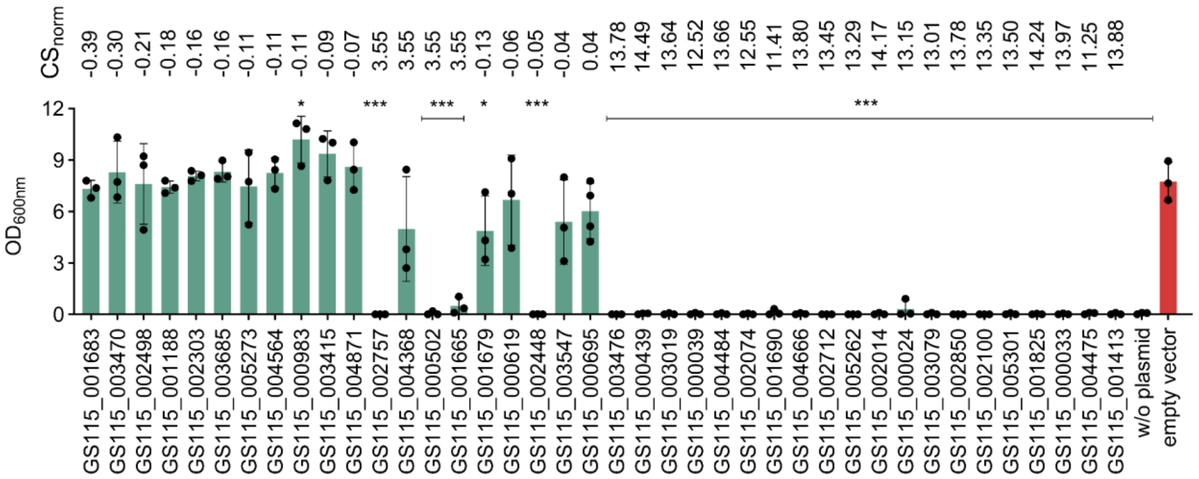


**Supplementary Figure 2** - Experimental CS validations obtained from CRISPR-Cas9 screen. Final $OD_{600}$ of Cas9-expressing NHEJ-deficient cells expressing 24 high CS and 16 low CS sgRNAs. Transformants were grown in SD-H for two days right after electroporation, followed by subculturing in fresh SD-H media and were allowed to grow for three more days. Cells were also transformed with an empty vector as the control to show the impact of the presence of sgRNA. GS115 containing no plasmid was used as the negative control, showing no growth post electroporation. Validation experiments were done in three biological replicates. The bars show the mean of the replicates, data points represent OD of each individual replicate, and the error bars represent one standard deviation. * $p < 0.05$, *** $p < 0.0005$ ; one-tailed unpaired t-test)

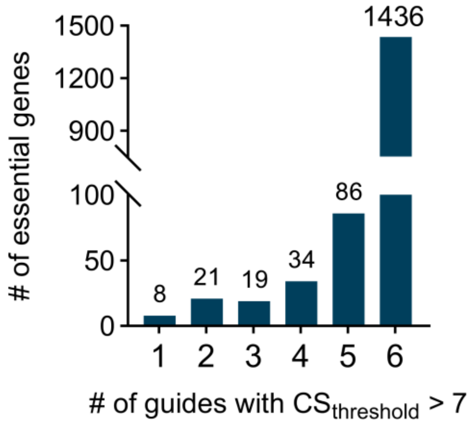


**Supplementary Figure 3** - Predicted essential gene coverage with activity-validated sgRNAs with a $CS_{threshold}$ > 7. More than 98% of the predicted essential genes are covered with more than one highly active sgRNA further validating the essential gene identification.

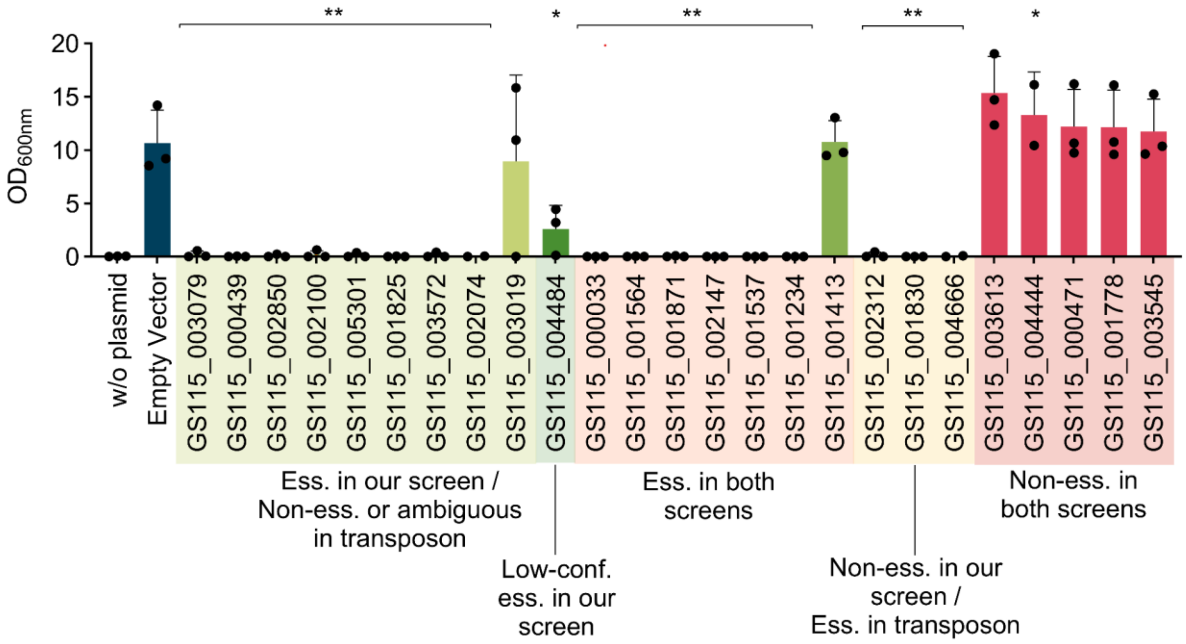


**Supplementary Figure 4 -** Experimental validation of CRISPR-Cas9 essential and non-essential genes from acCRISPR analysis. Final OD of Cas9 expressing cells expressing sgRNAs targeting essential, low-confidence essential, and non-essential genes. Transformants were grown in SD-H for two days right after electroporation, followed by subculturing in fresh SD-H media and were allowed to grow for three more days. Cells were also transformed with an empty vector as the control to show the impact of gene essentiality on cell fitness. GS115 containing no plasmid was used as the negative control, showing no growth post electroporation. Validation experiments were done in three biological replicates. The bars show the mean of the replicates, data points represent OD of each individual replicate, and the error bars represent one standard deviation. Statistical analysis is done between transformants with sgRNA-containing plasmids and the empty vector. * $p < 0.05$, ** $p < 0.005$ ; one-tailed unpaired t-test)

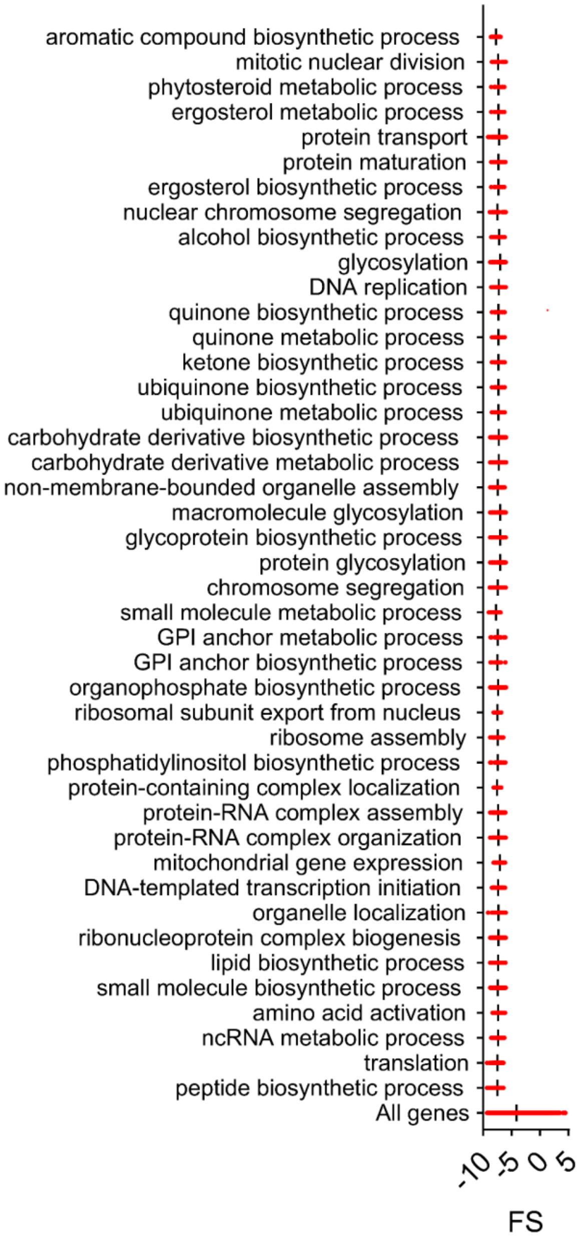


**Supplementary Figure 5** - Enriched GO biological process terms (adjusted p-value < 0.05) with the respective FS obtained from our CRISPR-Cas9 screen for identified essential genes associated to each term (Carlson and Pages, 2020; Wu et al., 2021; Yu et al., 2012). The FS values for each GO term were found to be significantly lower than those of all genes by unpaired t-test ($p < 0.0001$).

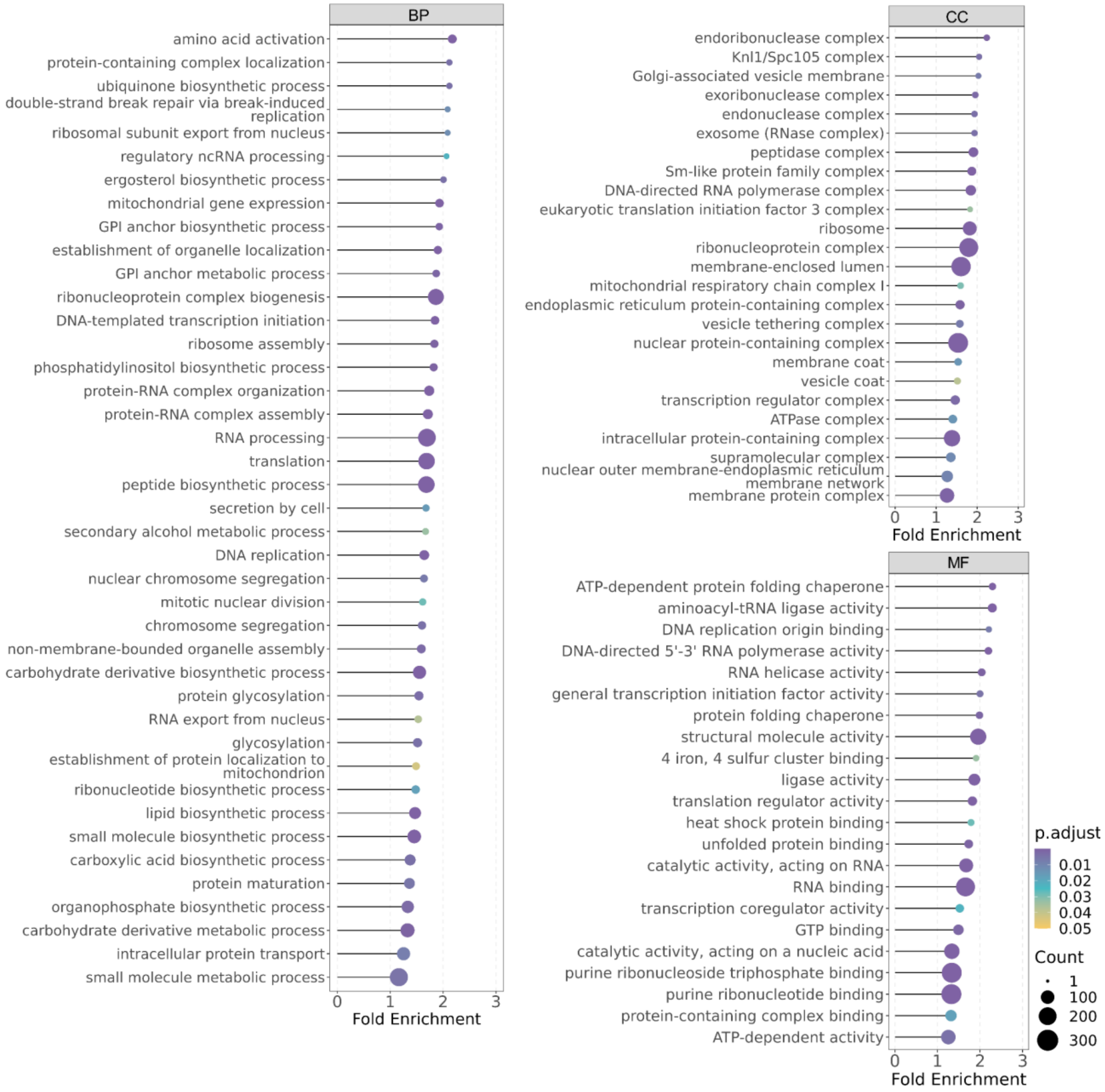


**Supplementary Figure 6. GO enrichment analysis for the consensus set of essential genes** Over-representation analyses were conducted by ClusterProfiler, and only the GO terms highly enriched (adjusted p-value < 0.05) in each category (BP: biological process, MF: molecular function, and CC: cellular component) were presented. (Carlson and Pages, 2020; Wu et al., 2021; Yu et al., 2012). The count represents the number of genes annotated in a specific term, and fold enrichment is defined as the ratio of the frequency of genes belonging to a specific enriched term in the gene set to the frequency of genes belonging to that term in the genome.

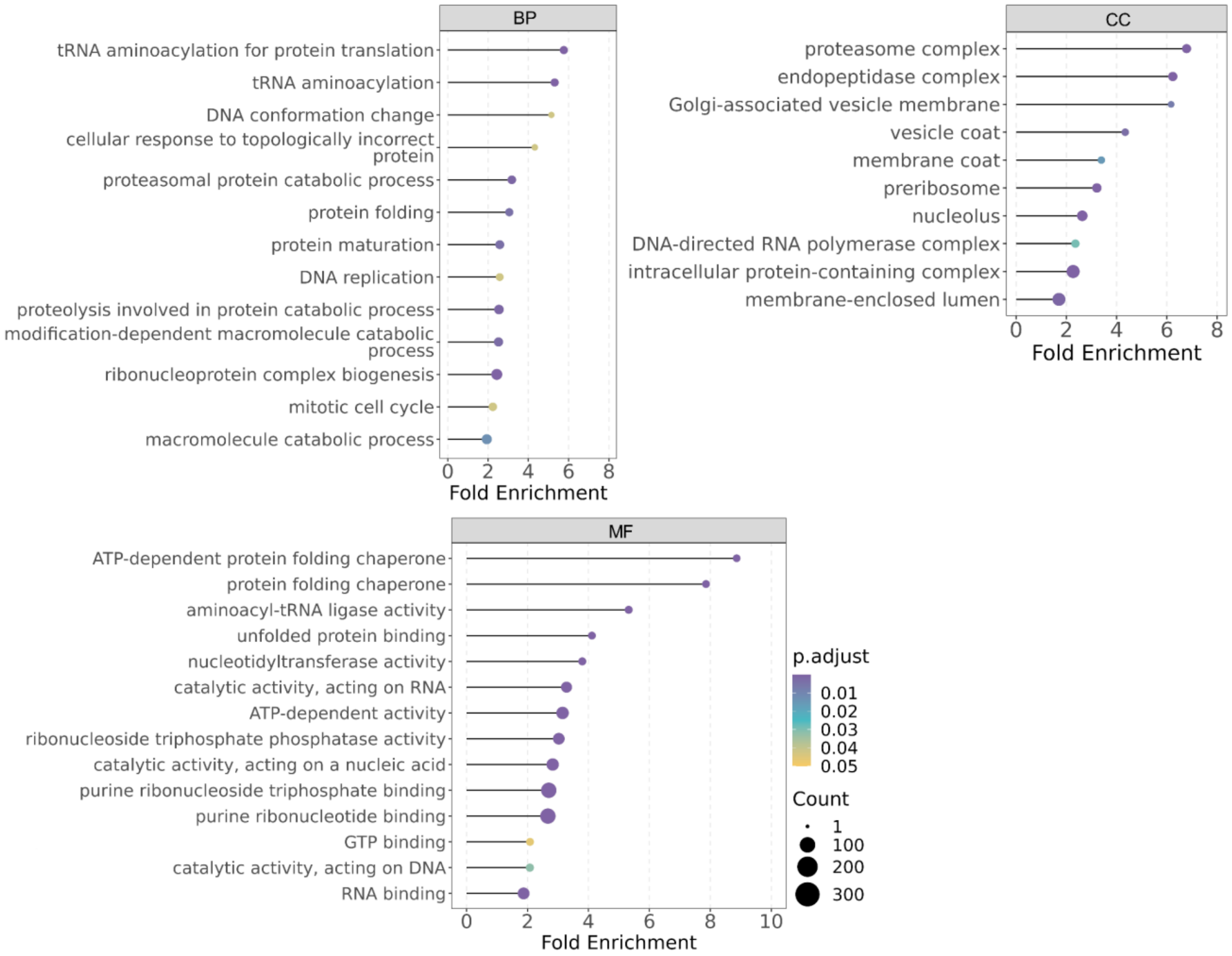


**Supplementary Figure 7. GO enrichment analysis for the yeast core set of essential genes** Over-representation analyses were conducted by ClusterProfiler, and only the GO terms highly enriched (adjusted p-value < 0.05) in each category (BP: biological process, MF: molecular function, and CC: cellular component) were presented. (Carlson and Pages, 2020; Wu et al., 2021; Yu et al., 2012). The count represents the number of genes annotated in a specific term, and fold enrichment is defined as the ratio of the frequency of genes belonging to a specific enriched term in the gene set to the frequency of genes belonging to that term in the genome.

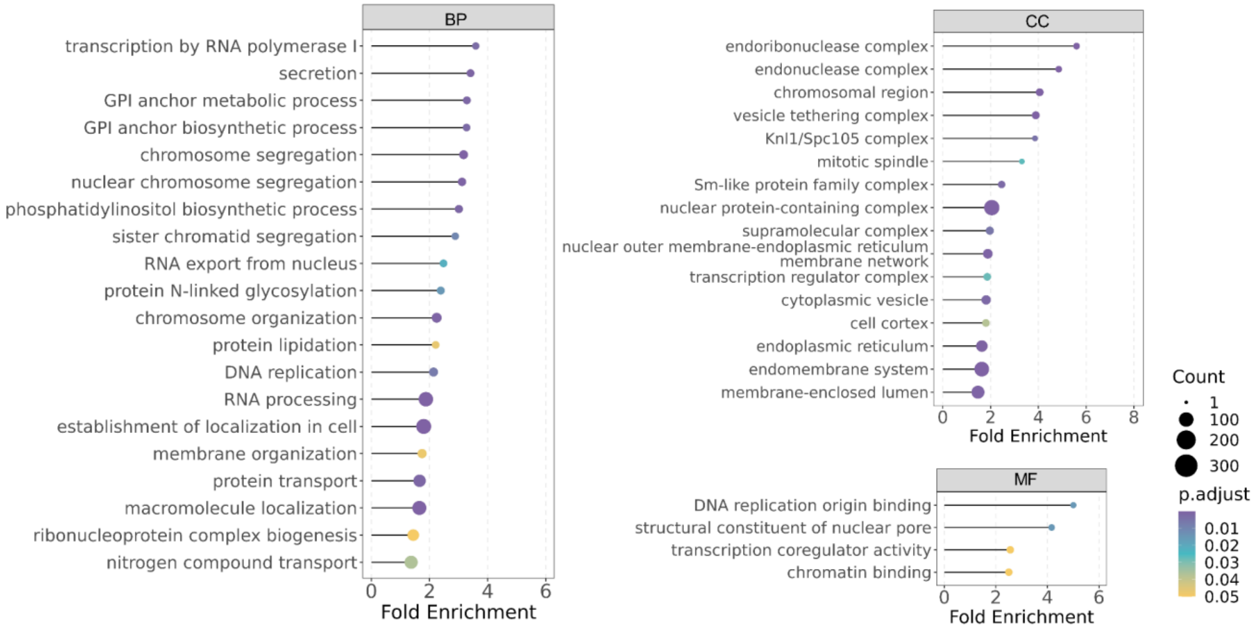


**Supplementary Figure 8. GO enrichment analysis for the *K. phaffii* specific set of essential genes** Over-representation analyses were conducted by ClusterProfiler, and only the GO terms highly enriched (adjusted p-value < 0.05) in each category (BP: biological process, MF: molecular function, and CC: cellular component) were presented. (Carlson and Pages, 2020; Wu et al., 2021; Yu et al., 2012). The count represents the number of genes annotated in a specific term, and fold enrichment is defined as the ratio of the frequency of genes belonging to a specific enriched term in the gene set to the frequency of genes belonging to that term in the genome.

**Supplementary Table 1**- Fold-coverage of the designed library for *K. phaffii* GS115 (Labun et al., 2019). More than 98% of the genes in the genome are designed to be targeted by six sgRNAs. Only seventeen genes lacked any designed sgRNAs due to redundancy in their sequence which led to the absence of unique guides.

| sgRNA fold-coverage | No. of genes with fold-coverage | % of genes with fold-coverage |
|---|---|---|
| 6 | 5132 | 94.58 |
| 5 | 37 | 0.68 |
| 4 | 71 | 1.31 |
| 3 | 70 | 1.29 |
| 2 | 64 | 1.18 |
| 1 | 35 | 0.64 |
| 0 | 17 | 0.31 |

**Supplementary Table 2 -** Strains used in this study.

| Strain | Genotype | Reference |
|---|---|---|
| E. coli TOP10 | F- mcrA Δ(mrr-hsdRMS-mcrBC) φ80lacZΔM15 ΔlacX74 recA1 araD139 Δ(ara-leu) 7697 galU galK rpsL (StrR) endA1 nupG λ | Thermo Fisher Scientific |
| GS115 | *his4* | Invitrogen |
| GS115 *his4*::*CAS9* | *his4*::*CAS9* | This study |
| GS115 *ku70* | *his4, ku70* | This study |
| GS115 *his4*::*CAS9 ku70* | *his4*::*CAS9, ku70* | This study |

**Supplementary Table 3 -** Plasmids used in this study.

| Plasmid | Description | Reference |
|---|---|---|
| D-227 | Contains Cas9 and sgRNA expression cassettes in *K. phaffii* | (Dalvie et al., 2020) |
| pCRISPRpp | sgRNA library backbone, contains PpHIS4 gene, PARS1 and AmpR | This study |
| pCRISPRyl | CRISPR/Cas9 vector for Yarrowia lipolytica, with AvrII site for sgRNA insertion | Addgene #70007 |
| BB3cK_pGAP_23*_pTEF_Cas9 | hCas9 under control of Tef1 for direct cloning of HH-sgRNA-HDV PCR products and episomal expression in P. pastoris and G418 selection | Addgene #104909 (Gassler et al., 2019) |
| pMJA089 | Expresses human Lysozyme in Pichia pastoris | Addgene #128518 (Yang et al., 2014) |

**Supplementary Table 4 -** Primers used in this study.

| Primer name | Primer sequence | Use |
|---|---|---|
| UDA.F | ATCAACGCTGTTCAACAAAATCTCAACA | *HIS4* loci upstream donor arm amplification |
| UDA.R | TGCACATTAACTTGAAGCTCAGTCG | *HIS4* loci upstream donor arm amplification |
| DDA.F | AGGACCAATTTGAGGAGCTGAT | *HIS4* loci downstream donor arm amplification |
| DDA.R | TGTTCCTTGGTGTATCCTGGCT | *HIS4* loci downstream donor arm amplification |
| D227_UDA.F | GAGCTTCAAGTTAATGTGCAATTAAGTA ATAGCAAGGTAAAGTAATACAGGGAGT | D-227 vector linearization for upstream donor arm integration |
| D227_UDA.R | GATTTTGTTGAACAGCGTTGATGACTCCC AAGTCTAAGGACTTGA | D-227 vector linearization for upstream donor arm integration |
| D227_DDA.F | GCCAGGATACACCAAGGAACAACCATGG GATATGTTTCACGTTTTGT | D-227 vector linearization for downstream donor arm integration |
| D227_DDA.R | CAGCTCCTCAAATTGGTCCTATGAAAGA GTGAGAGGAAAGTACCTG | D-227 vector linearization for downstream donor arm integration |
| HIS4_ID.F | CTAAACGAAAGACTACATTTCTAGATGA GTTTGCC | PCR amplification of the *HIS4* loci to check for Cas9 integration |
| HIS4_ID.R | CCTGACGTTATCTATAGAGAGATCAATG GCTC | PCR amplification of the *HIS4* loci to check for Cas9 integration |
| Ori_AmpR.F | AATACGGTTATCCACAGAATCAGGGGA | Ori_AmpR PCR amplification from pCRISPRyl |
| Ori_AmpR.R | GGAAATGTGtGCGGAACC | Ori_AmpR PCR amplification from pCRISPRyl |
| PARS1.F | ACGAGGCCCAGATCCTCTATTAATTAACC TAGGGGTACCTTCAAGTTTCGTTAAGCAG GA | PARS1 PCR amplification from the genome |
| PARS1.R | CCTGATTCTGTGGATAACCGTATTACTAG TGATTGATATTGGAACCTGCTGTCATT | PARS1 PCR amplification from the genome |
| pTEF1.F | AATAGGGGTTCCGCGCACATTTCCGCATG Cgcatcaccatctgaatatttgaccgct | TEF1 promoter amplification from BB3cK_pGAP_23*_pTEF_Cas9 plasmid |
| pTEF1.R | gcaggtagcaagggaaatgtcatGGTTACCgaccgccctta gattagattgctatgc | TEF1 promoter amplification from BB3cK_pGAP_23*_pTEF_Cas9 plasmid |
| CYC1t.F | gaaactgggacttatttaaGGGCCCtcatgtaattagttatgtca cgcttac | CYC1 terminator amplification from BB3cK_pGAP_23*_pTEF_Cas9 plasmid |

| | | |
|---|---|---|
| CYC1t.R | ATTCAAGCTAATATGGCTGATGATCCTCTAACCTACTACGcatgaattagcgccagcttg | CYC1 terminator amplification from BB3cK_pGAP_23*_pTEF_Cas9 plasmid |
| ptRNA1_tRNA1.F | AGCCATATTAGCTTGAATGATTGGATTTTTTGTAGCTTTATAAGCAGCTTTTTCTTGAAG | gRNA expression cassette amplification from D-227 |
| ptRNA1_tRNA1.R | GGTTAATGTCATGATAATAATGGTTTCTTAgacgtAAAAAAAAGCACCGACTCGGTGCC | gRNA expression cassette amplification from D-227 |
| PpHIS4.F | atgacatttcccttgctacctg | Pichia pastoris HIS4 gene from pIB1 |
| PpHIS4.R | aagcgtgacataactaattacatgaGGGCCCttaaataagtcccagtttctccatacg | Pichia pastoris HIS4 gene from pIB1 |
| 5'_60mer | tagtggtagaaccaccgcttgtcgcgcggtagaccggggttcaattccccgtcgcggagc | 5' 60mer linker added to the designed sgRNAs in the library |
| 3'_60mer | gttttagagctagaaatagcaagttaaaataaggctagtccgttatcaacttgaaaaagt | 3' 60mer linker added to the designed sgRNAs in the library |
| qPCR_GW.F | GCGCCTTATCCGGTAACTATC | qPCR experiment primer to count the copy number of miniprepped library from GS115 cells |
| qPCR_GW.R | CTACATACCTCGCTCTGCTAATC | qPCR experiment primer to count the copy number of miniprepped library from GS115 cells |
| NGS1.F | AATGATACGGCGACCACCGAGATCTACACTCTTTCCCTACACGACGCTCTTCCGATCTAGtgtagaccggggttcaattccc | Forward Illumina primer used for NGS |
| NGS2.F | AATGATACGGCGACCACCGAGATCTACACTCTTTCCCTACACGACGCTCTTCCGATCTGTagtgtagaccggggttcaattccc | Forward Illumina primer used for NGS |
| NGS3.F | AATGATACGGCGACCACCGAGATCTACACTCTTTCCCTACACGACGCTCTTCCGATCTCAGTagtgtagaccggggttcaattccc | Forward Illumina primer used for NGS |
| NGS4.F | AATGATACGGCGACCACCGAGATCTACACTCTTTCCCTACACGACGCTCTTCCGATCTTCCAGTagtgtagaccggggttcaattccc | Forward Illumina primer used for NGS |
| NGS1.R | CAAGCAGAAGACGGCATACGAGATTCGCCTTGGTGACTGGAGTTCAGACGTGTGCTCTTCCGATCTAAGTTGATAACGGACTAGCCT | Reverse Illumina primer used for NGS |
| NGS2.R | CAAGCAGAAGACGGCATACGAGATATAGCGTCGTGACTGGAGTTCAGACGTGTGCTCTTCCGATCTAAGTTGATAACGGACTAGCCT | Reverse Illumina primer used for NGS |
| NGS3.R | CAAGCAGAAGACGGCATACGAGATGAAGAAGTGTGACTGGAGTTCAGACGTGTGCTCTTCCGATCTAAGTTGATAACGGACTAGCCT | Reverse Illumina primer used for NGS |
| NGS4.R | CAAGCAGAAGACGGCATACGAGATATTCTAGGGTGACTGGAGTTCAGACGTGTGCT | Reverse Illumina primer used for NGS |

| | | |
|---|---|---|
| | CTTCCGATCTAAGTTGATAACGGACTAGCCT | |
| NGS5.R | CAAGCAGAAGACGGCATACGAGATCGTTACCAGTGACTGGAGTTCAGACGTGTGCTCTTCCGATCTAAGTTGATAACGGACTAGCCT | Reverse Illumina primer used for NGS |
| NGS6.R | CAAGCAGAAGACGGCATACGAGATGTCTGATGGTGACTGGAGTTCAGACGTGTGCTCTTCCGATCTAAGTTGATAACGGACTAGCCT | Reverse Illumina primer used for NGS |
| NGS7.R | CAAGCAGAAGACGGCATACGAGATTTACGCACGTGACTGGAGTTCAGACGTGTGCTCTTCCGATCTAAGTTGATAACGGACTAGCCT | Reverse Illumina primer used for NGS |
| NGS8.R | CAAGCAGAAGACGGCATACGAGATTTGAATAGGTGACTGGAGTTCAGACGTGTGCTCTTCCGATCTAAGTTGATAACGGACTAGCCT | Reverse Illumina primer used for NGS |
| NGS9.R | CAAGCAGAAGACGGCATACGAGATTGCTGAGCGTGACTGGAGTTCAGACGTGTGCTCTTCCGATCTAAGTTGATAACGGACTAGCCT | Reverse Illumina primer used for NGS |

**Supplementary Table 5 -** sgRNAs used in this study.

| gRNA sequence | Targeting gene |
|---|---|
| AGTTGAGTCATCCGTTACAG | *HIS4* |
| AAGCAATACGACATCCACGA | *KU70* |
| GAAGCTCAAGGAATCAAGAA | GS115_000439_6.0 |
| AAGCACTCAGACTCTCCCTT | GS115_003019_6.0 |
| AAACATCATTATCGGGCGTG | GS115_000695_1.0 |
| ATGTCGCTCTTCTTCATAGA | GS115_001683_4.0 |
| TGGAGGTCTAATCATAACTG | GS115_003470_2.0 |
| CGGAGAGTTACAATTCTCCG | GS115_002498_2.0 |
| GGCAATATGCCATTGCCACC | GS115_001188_2.0 |
| CTTTGTGGCCTCTAAATGAG | GS115_002303_4.0 |
| ACATCCTGCTGCAAAGATGT | GS115_003685_4.0 |
| GTCCACCTCCAAATCCACCA | GS115_005273_3.0 |
| GGACTCAGATCCAGACTCGG | GS115_004564_1.0 |
| CCAAAGGATCCCAGGCGCTA | GS115_000983_5.0 |
| AAGCAATACGACATCCACGA | GS115_003415_5.0 |
| CCAGAGAATCAAATTGCCAA | GS115_004871_6.0 |
| ATATAGGGATCTTCCAAAGG | GS115_002757_4.0 |
| GCAGAAGCGCTCCAGTACGA | GS115_004368_2.0 |
| GAGCTTCAGCAGTGGTACAA | GS115_000502_5.0 |
| GCCACCCAAATCCACCCTCG | GS115_001665_3.0 |
| CTATGAAACAAAAACGTTCT | GS115_001679_6.0 |
| CTCATTCAAAGAATTGAATG | GS115_000619_4.0 |
| AATTCCTTCAACAAATCCGG | GS115_002448_5.0 |
| TCCAATGACATGAAAGCCAT | GS115_003547_2.0 |
| TCACGTCGACTGATGCGCAT | GS115_003476_1.0 |
| GAAGCTCAAGGAATCAAGAA | GS115_000439_6.0 |
| AAGCACTCAGACTCTCCCTT | GS115_003019_6.0 |
| TGGCGAACTAATCTCCGTAC | GS115_000039_3.0 |
| GGTGTCTCTTAGCTTCTCAG | GS115_004484_4.0 |
| TACTGTTTACAAGGGAACGA | GS115_002074_3.0 |
| TGAGATGGCTGAGCGGAGGG | GS115_001690_3.0 |
| GATGAAAATGAGATTCTGGC | GS115_004666_5.0 |
| TACAGCAGCCATCTTGTAAG | GS115_002712_5.0 |
| CATAGTGAATCAGTCAACCT | GS115_005262_1.0 |
| GTGCTTTCAACATAATCTGG | GS115_002014_1.0 |
| GGTGAAACAAGAGGTTGCGA | GS115_000024_2.0 |
| ATCTGGTAGTGGAGAACCGG | GS115_003079_4.0 |

| | |
|---|---|
| ACTTGCTGAACAAATCGGGT | GS115_002850_6.0 |
| GCTTCTATACAAATCGGATG | GS115_002100_6.0 |
| GTTGAGACTGTCAGAAATTG | GS115_005301_5.0 |
| GCTCAGGACAGATAACGTGC | GS115_001825_5.0 |
| GTTCACAGAGGCCTACAGAC | GS115_000033_6.0 |
| AAGAAACATTAACACAACAT | GS115_004475_2.0 |
| GTGAGATTTGAGATTCAAGG | GS115_001413_2.0 |
| ACCAACGGTAAAGTTCCTGA | GS115_003572_4.0 |
| GTGGCCAAGAGAACTCCAGC | GS115_001564_4.0 |
| GTCCTCACGATCGTTGACAA | GS115_001871_2.0 |
| TGGTGTAACAAATATCGCCT | GS115_002147_5.0 |
| TGCCAAACGAAGAATCAACC | GS115_001537_3.0 |
| TGCAACCATTGAGTAGTAGT | GS115_002312_4.0 |
| GGAGCTCCAGTGGGGACAGT | GS115_001830_4.0 |
| ACTATGACTCGGAAGAAAGA | GS115_001234_2.0 |
| AGACACACAGGATAGCGGGC | GS115_003613_4.0 |
| TCTACGACGCCGTTTAGCAC | GS115_004444_4.0 |
| ACCGAAGAAGATGATGCGGA | GS115_000471_2.0 |
| GTAAAAACCCTATCAGGCCG | GS115_001778_2.0 |
| TCCAACTCCAAGGATACCAT | GS115_003545_6.0 |

**Supplementary Table 6 -** Comprehensive breakdown of quality scores used to design the final library.

| Quality score | Criteria |
|---|---|
| 1 | Seed_MM0 = 0, MM0 = 0, Self_complementarity = 0, MM1 = 0, MM2 = 0, MM3 = 0 |
| 2 | Seed_MM0 = 0, MM0 = 0, Self_complementarity = 0, MM1 = 0, MM2 = 0, MM3 = 1 |
| 3 | Seed_MM0 = 0, MM0 = 0, Self_complementarity = 0, MM1 = 0, MM2 = 0, MM3 = 2 |
| 4 | Seed_MM0 = 0, MM0 = 0, Self_complementarity = 0, MM1 = 0, MM2 = 0, MM3 = 3 |
| 5 | Seed_MM0 = 0, MM0 = 0, Self_complementarity = 0, MM1 = 0, MM2 = 1, MM3 = 0 |
| 6 | Seed_MM0 = 0, MM0 = 0, Self_complementarity = 0, MM1 = 0, MM2 = 1, MM3 = 1 |
| 7 | Seed_MM0 = 0, MM0 = 0, Self_complementarity = 0, MM1 = 0, MM2 = 1, MM3 = 2 |
| 8 | Seed_MM0 = 0, MM0 = 0, Self_complementarity = 0, MM1 = 0, MM2 = 1, MM3 = 3 |
| 9 | Seed_MM0 = 0, MM0 = 0, Self_complementarity = 0, MM1 = 0, MM2 = 2, MM3 = 0 |
| 10 | Seed_MM0 = 0, MM0 = 0, Self_complementarity = 0, MM1 = 0, MM2 = 2, MM3 = 1 |
| 11 | Seed_MM0 = 0, MM0 = 0, Self_complementarity = 0, MM1 = 0, MM2 = 2, MM3 = 2 |
| 12 | Seed_MM0 = 0, MM0 = 0, Self_complementarity = 0, MM1 = 0, MM2 = 2, MM3 = 3 |
| 13 | Seed_MM0 = 1, MM0 = 1 |

**Supplementary Table 7 -**Number of sgRNAs in the designed library with their respective quality score.

| Quality Score | No. of gRNAs | % of the library |
|---|---|---|
| 1 | 31433 | 99.36 |
| 2 | 61 | 0.19 |
| 3 | 22 | 0.07 |
| 4 | 5 | 0.02 |
| 5 | 18 | 0.06 |
| 6 | 6 | 0.02 |
| 7 | 1 | 0.003 |
| 8 | 1 | 0.003 |
| 9 | 0 | 0 |
| 10 | 0 | 0 |
| 11 | 2 | 0.006 |
| 12 | 1 | 0.003 |
| 13 | 84 | 0.26 |
| Non-targeting | 350 | 1.089155 |
| Total | 32485 | 100 |

**Supplementary Table 8 -** Transformation efficiencies measured as ×$10^6$ transformants for 10 pooled transformations, for all replicates in the control and treatment strains.

| | Transformation efficiency (×$10^6$) | | |
|---|---|---|---|
| Strain | Replicate 1 | Replicate 2 | Replicate 3 |
| GS115 | 7.03 | 10.93 | 6.3 |
| GS115 Cas9 | 8.35 | 6.85 | 10.7 |
| GS115 Δku70 | 5.1 | 5.4 | 8.86 |
| GS115 Δku70 Cas9 | 4.9 | 7.55 | 9.55 |

**Supplementary Table 9 -** Library plasmid copy number and fold-coverage from miniprepped samples for all strains and replicates.

| Strain-Replicate-Day | Plasmid copy number | Fold-coverage |
|---|---|---|
| GS115-Rep1-Day3 | 3.21E+08 | 9441 |
| GS115-Rep1-Day6 | 4.40E+08 | 12941 |
| GS115-Rep2-Day3 | 1.36E+09 | 40000 |
| GS115-Rep2-Day6 | 1.98E+08 | 5824 |
| GS115-Rep3-Day3 | 1.32E+09 | 38824 |
| GS115-Rep3-Day6 | 1.04E+08 | 3059 |
| GS115 Cas9-Rep1-Day3 | 4.08E+08 | 12000 |
| GS115 Cas9-Rep1-Day6 | 1.50E+08 | 4412 |
| GS115 Cas9-Rep2-Day3 | 1.10E+09 | 32353 |
| GS115 Cas9-Rep2-Day6 | 1.93E+08 | 5676 |
| GS115 Cas9-Rep3-Day3 | 6.62E+08 | 19471 |
| GS115 Cas9-Rep3-Day6 | 1.43E+08 | 4206 |
| GS115 Δku70-Rep1-Day3 | 5.72E+07 | 1683 |
| GS115 Δku70-Rep1-Day6 | 1.40E+08 | 4129 |
| GS115 Δku70-Rep2-Day3 | 1.63E+08 | 4803 |
| GS115 Δku70-Rep2-Day6 | 5.28E+07 | 1552 |
| GS115 Δku70-Rep3-Day3 | 1.74E+08 | 5115 |
| GS115 Δku70-Rep3-Day6 | 3.72E+07 | 1095 |
| GS115 Δku70 Cas9-Rep1-Day3 | 1.89E+08 | 5562 |
| GS115 Δku70 Cas9-Rep1-Day6 | 8.90E+07 | 2616 |
| GS115 Δku70 Cas9-Rep2-Day3 | 2.69E+08 | 7918 |
| GS115 Δku70 Cas9-Rep2-Day6 | 1.69E+08 | 4974 |
| GS115 Δku70 Cas9-Rep3-Day3 | 1.19E+08 | 3497 |
| GS115 Δku70 Cas9-Rep3-Day6 | 1.48E+08 | 4344 |

| Strain | Time | Replicate | Pearson r |
|---|---|---|---|
| GS115 | Day3 | 1 vs. 2 | 0.89 |
| | | 1 vs. 3 | 0.8874 |
| | | 2 vs. 3 | 0.8911 |
| GS115 | Day6 | 1 vs. 2 | 0.8236 |
| | | 1 vs. 3 | 0.8216 |
| | | 2 vs. 3 | 0.8453 |
| GS115 Cas9 | Day3 | 1 vs. 2 | 0.9829 |
| | | 1 vs. 3 | 0.9852 |
| | | 2 vs. 3 | 0.9823 |
| GS115 Cas9 | Day6 | 1 vs. 2 | 0.9792 |
| | | 1 vs. 3 | 0.98 |
| | | 2 vs. 3 | 0.9821 |
| GS115 Δku70 | Day3 | 1 vs. 2 | 0.8759 |
| | | 1 vs. 3 | 0.8859 |
| | | 2 vs. 3 | 0.8962 |
| GS115 Δku70 | Day6 | 1 vs. 2 | 0.6654 |
| | | 1 vs. 3 | 0.6936 |
| | | 2 vs. 3 | 0.8015 |
| GS115 Δku70 Cas9 | Day3 | 1 vs. 2 | 0.9799 |
| | | 1 vs. 3 | 0.9836 |
| | | 2 vs. 3 | 0.9844 |
| GS115 Δku70 Cas9 | Day6 | 1 vs. 2 | 0.9522 |
| | | 1 vs. 3 | 0.9745 |
| | | 2 vs. 3 | 0.9531 |

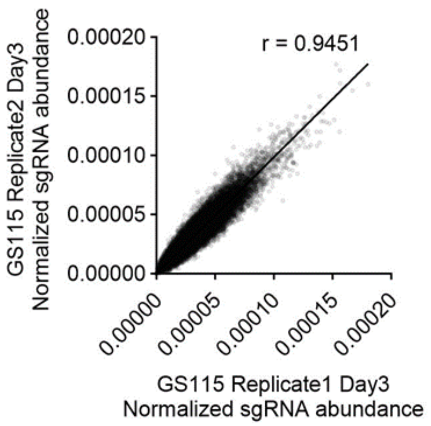


**Supplementary Table 10 -** Correlation of normalized sgRNA abundance between biological replicates. The plot shows the normalized sgRNA abundance between replicate 1 and 2 for GS115. Linear regression between replicate 1 and 2 yields a Pearson coefficient of 0.9451. The table represents the correlation of normalized sgRNA abundance for all strains and all possible combinations of replicates.

#### References


Carlson, M., Pages, H., 2020. AnnotationForge: tools for building SQLite-based annotation data packages. R Packag. version 1.32. 0.

Dalvie, N.C., Leal, J., Whittaker, C.A., Yang, Y., Brady, J.R., Love, K.R., Love, J.C., 2020. Host-Informed Expression of CRISPR Guide RNA for Genomic Engineering in Komagataella phaffii. ACS Synth. Biol. 9, 26–35.

Gassler, T., Heistinger, L., Mattanovich, D., Gasser, B., Prielhofer, R., 2019. CRISPR/Cas9-Mediated Homology-Directed Genome Editing in Pichia pastoris. Methods Mol. Biol. 1923, 211–225.

Labun, K., Montague, T.G., Krause, M., Torres Cleuren, Y.N., Tjeldnes, H., Valen, E., 2019. CHOPCHOP v3: expanding the CRISPR web toolbox beyond genome editing. Nucleic Acids Res. 47, W171–W174.

Wu, T., Hu, E., Xu, S., Chen, M., Guo, P., Dai, Z., Feng, T., Zhou, L., Tang, W., Zhan, L., Fu, X., Liu, S., Bo, X., Yu, G., 2021. clusterProfiler 4.0: A universal enrichment tool for interpreting omics data. Innovation (Camb) 2, 100141.

Yang, J., Nie, L., Chen, B., Liu, Y., Kong, Y., Wang, H., Diao, L., 2014. Hygromycin-resistance vectors for gene expression in Pichia pastoris. Yeast 31, 115–125.

Yu, G., Wang, L.-G., Han, Y., He, Q.-Y., 2012. clusterProfiler: an R package for comparing biological themes among gene clusters. OMICS 16, 284–287.
